# Integrated Stress Response Triggered by Excessive Glycosylation Drives Thoracic Aortic aneurysm

**DOI:** 10.1101/2024.05.31.596791

**Authors:** Antonio Rochano-Ortiz, Irene San Sebastian-Jaraba, Carmen Zamora, Carolina Simó, Virginia García-Cañas, Sacramento Martínez-Albaladejo, María José Fernandez-Gomez, Tiago R. Velho, María Jesús Ruíz-Rodríguez, Amanda Leal-Zafra, Enrique Gabandé, Sara Martinez-Martinez, Andrea Guala, Óscar Lorenzo, Luis Miguel Blanco-Colio, José Luís Martín-Ventura, Gisela Teixido-Tura, Alberto Forteza, J. Francisco Nistal, Juan Miguel Redondo, Nerea Méndez-Barbero, María Mittelbrunn, Jorge Oller

## Abstract

Thoracic aortic aneurysms and dissections (TAAD) are marked by degenerative changes in the aortic media. Marfan syndrome is the most common inherited connective tissue disorder associated with TAAD. While vascular smooth muscle cell (VSMC) metabolism is emerging as a targetable driver of aortic aneurysm, surgical interventions remain the primary strategy to prevent aortic dissection. Our research indicates that the hexosamine biosynthetic pathway (HBP), a branch of glycolysis, is upregulated in aortas from the *Fbn1^C1041G/+^* Marfan Syndrome mouse model. Enhancing HBP activity promotes aortic dilation and accumulation glycan-rich extracellular matrix, contributing to aortic medial degeneration in wild-type mice. Mechanistically, fueling HBP activity induces VSMC dysfunction through excessive glycosylation, which activates the Integrated Stress Response (ISR). Pharmacological inhibition of HBP, along with ISR inhibition, successfully reverses aortic dilation and aortic medial degeneration in *Fbn1^C1041G/+^* Marfan Syndrome mouse model. Additionally, Marfan Syndrome patients show elevated levels of HBP metabolites in blood plasma and serum, and heightened HBP-ISR signaling in patients with TAAD. These findings unveil a potential causative role for the HBP-ISR axis in medial degeneration in human TAAD, underscoring the need for evaluating HBP and ISR pathway as novel biomarkers and therapeutic strategies for thoracic aortic aneurysm.

## Introduction

Thoracic aortic aneurysm and dissection (TAAD) represent substantial threats to health and longevity in developed nations, constituting a primary cause of morbidity and mortality^1^. This highlights the importance of developing pharmacological interventions to slow or halt aortic diameter growth and prevent aortic rupture. This condition evolves through a gradual dilation of the aorta, which is attributed to dysfunction in vascular smooth muscle cells (VSMCs) and extensive extracellular matrix remodeling^1,2^. Genetic factors play a pivotal role in TAAD, with approximately one in four cases having a known genetic basis^3^. One of the most prevalent inherited connective tissue disorders is the Marfan syndrome (MFS). MFS is considered a rare disorder (OMIM #154700), due to its documented incidence of 1 in 5000 individuals. MFS is primarily caused by dominant pathogenic variants in the gene *FBN1* encoding the extracellular protein Fibrillin1^4^. Individuals affected by MFS commonly exhibit elongated bones, lens luxation, and unfortunately, a shortened life expectancy, largely due to the complications arising from TAAD. Indeed, TAAD and subsequent rupture contribute to over 90% of deaths among MFS patients^5^. Currently, pharmacologic interventions for preventing or reversing TAAD remain elusive, leaving open surgery as the primary options for preventing vascular complications. It is therefore, impetrative to address this critical gap in treatment options is imperative to improve the prognosis and quality of life for patients affected by TAAD^5,6^.

VSMCs constitute the predominant cell type within the medial layer of the aorta and play a crucial role in regulating arterial lumen size by directly modulating vascular wall contraction. VSMCs possess the remarkable ability to transition reversibly between pathological, or secretory, and quiescent phenotypes. The secretory phenotype is characterized by increased accumulation and remodeling of the extracellular matrix, thereby altering the mechanical properties of the aortic wall and promoting aneurysm formation^7–9^.

Aortic medial degeneration, previously referred to as cystic or medial necrosis degeneration, stands as a histological hallmark of TAAD^10^. Initially described by Babes and Mironescu in 1910, recently acquired significant attention^11–14^. Aortic medial degeneration is typified by the fragmentation of elastic fibers, fibrosis, excessive deposition of ground-mucoid matrix rich in proteoglycans (PGs) and glycosaminoglycans (GAGs). The increased presence of PGs and GAGs is suggested to contribute to heightened stiffness and mechanical stress through their osmotic activity within the aortic wall. Broad research has established associations between aortic medial degeneration and hereditary vascular disorders, as well as vascular aging. Aortic medial degeneration is notably prevalent in TAAD-related diseases such as MFS, Loeys-Dietz and Ehlers-Danlos syndromes; and annuloaortic ectasia, often occurring due to degenerative changes in the aortic wall, particularly in elderly patients^15–17^. Despite its significance, the molecular mechanisms underlying the accumulation of glycan-rich matrix in aortic medial degeneration and its impact on aortic homeostasis remain poorly understood^18–20^.

Previously, we have demonstrated that vascular metabolism and plays an essential role in the development of both, atherosclerotic and MFS aortic aneurysms and subsequent rupture^21,22^. The Hexosamine Biosynthetic Pathway (HBP), branching off from glycolysis, is responsible for generating uridine diphosphate N-acetylglucosamine (UDP-GlcNAc), a critical substrate for assembling GAGs and PGs, as well as for post-translational protein modification via O-linked β-N-acetylglucosamine (O-GlcNac)^23,24^. Recent research has begun to uncover the multifaceted role of HBP in various pathological conditions, including cancer metastasis and heart failure and hypertrophy^25–28^. Accumulation of PGs, GAGs, and glycosylated proteins mediated by the HBP, triggers the activation of the Integrated Stress Response (ISR) by Endoplasmatic Reticulum stress activation^29,30^. While short-term ISR primarily promotes cell survival, prolonged exposure to chronic stressors leads to maladaptive signaling pathways, which ultimately resulting in cellular dysfunction, apoptosis, and necrosis^31^.

Our study provides the first evidence indicating that aortic medial degeneration in TAAD is driven by excessive activity of the HBP. Increasing HBP activity through exogenous glucosamine administration or lentiviral vectors expressing Glutamine-Fructose-6-Phosphate Transaminase-2 (GFPT2), the rate-limiting HBP enzyme, induced aortic dilation and aortic medial degeneration. Conversely, treatment with an HBP inhibitor restores aortic homeostasis in a murine model of MFS by reducing glycosylation levels, aortic medial degeneration, and ISR signaling. Additionally, administration of an ISR inhibitor in a MFS mouse model, reversed TAAD, thereby improving aortic homeostasis. Importantly, both HBP and ISR were increased in aortic samples from MFS patients. Furthermore, GAGs and HBP metabolites were elevated in blood samples from both MFS mice and patients. These findings highlight the potential of HBP and ISR as novel biomarkers, mediators, and therapeutic targets for the management of aortic dilation and the prevention of aortic dissection in MFS and potentially other related conditions.

## Results

### Hexosamine biosynthetic pathway is increased in aortas from a murine model of Marfan Syndrome

To elucidate the molecular mechanisms underlying aortic medial degeneration, we analyzed our previous transcriptional data using from next-generation sequencing in *Fbn1^C1041G/+^*, a MFS-mice model^21^. Remarkably, key features observed in MFS patients, including syndromic phenotype, aortic dilatation, aneurysms, and aortic medial degeneration, are reproduced in *Fbn1^C1041G/+^* mice^32^. Aortas from 24-week-old *Fbn1^+/+^* and *Fbn1^C1041G/+^* mice were analyzed, showing an intermediate stage of TAAD. Our analysis using Ingenuity Pathway Analysis revealed a significant increase in the expression of genes related to UDP-N-acetyl-D-glucosamine Biosynthesis (also known as the Hexosamine Biosynthetic Pathway, HBP) within the aortas of MFS-mice (p-value= 0.047). Despite being a minor branch of glycolysis, constituting only 2–5% of total glucose metabolism under homeostatic conditions, the HBP plays a crucial role through its final metabolite, UDP-GlcNAc. This serves as substrate for glycosylation, a cornerstone for diverse cellular processes. Notably, UDP-GlcNAc is utilized for post-translational modification of intracellular proteins through O-GlcNAcylation (O-GlcNac) and also contributes to the synthesis and assembly of essential cellular components, including glycolipids, GAGs, PGs, and glycoproteins^23,24^(Fig. 1A,B).

**Figure 1:**
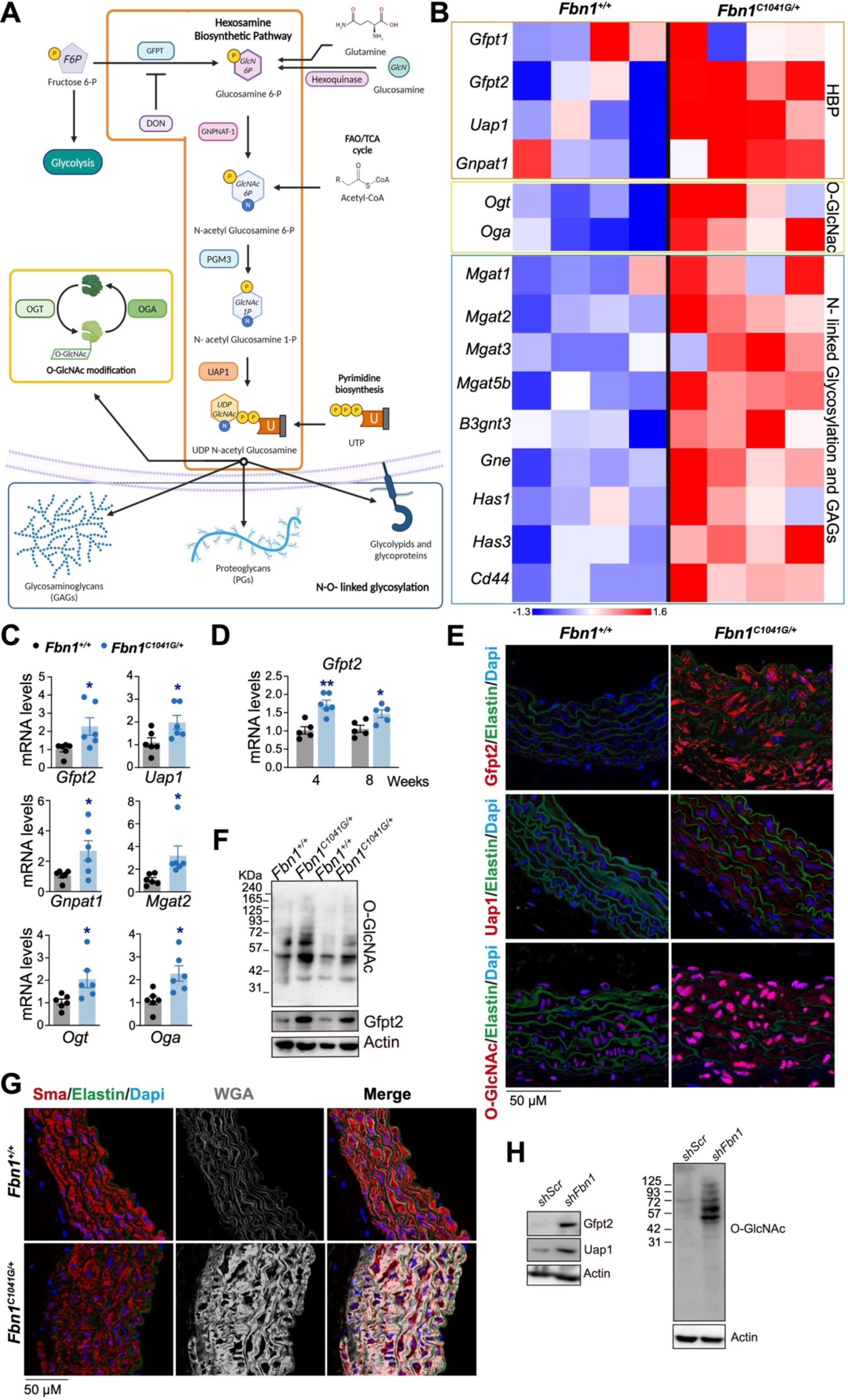
The HBP genes are upregulated in the aortas of a murine model MFS. (**A**) Scheme depicting the HBP role in O-glycosylation, N-linked glycosylation and GAGs. (**B**) Gene expression heatmap of HBP, O-glycosylation,N-linked glycosylation and GAGs-related genes from RNA-sequencing analysis of aortic medial tissue from 24 weeks old *Fbn1*^+/+^ and from *Fbn1^C1041G/+^* mice (n=4). (**C**) Quantitative reverse transcription polymerase chain reaction analysis of *Gfpt2, Uap1, Gnpat1, Mgat2, Ogt, Oga* mRNA relative expression in aortic extracts from 20/24-week-old *Fbn1^C1041G/+^* and *Fbn1*^+/+^ mice. (**D**) Quantitative reverse transcription polymerase chain reaction analysis of *Gfpt2* mRNA relative expression in aortic extracts from 4- or 8-weeks-old *Fbn1^C1041G/+^* and *Fbn1*^+/+^ mice. (**E**) Representative confocal imaging of Gfpt2, Uap1, O-GlcNac (red); elastin (green, autofluorescence), and DAPI–stained nuclei (blue) in the ascending aorta from 20-week-old *Fbn1^C1041G/+^* and *Fbn1*^+/+^ mice (n=6). (**F**) Representative immunoblot analysis of O-GlcNac proteins, Gfpt2 in aortic extracts from 20-week-old *Fbn1^C1041G/+^* and *Fbn1*^+/+^ mice (n=6). Actin was used as loading control. (**G**) Representative confocal imaging of WGA (gray), Smooth muscle actin (Sma, red), elastin (green, autofluorescence), and DAPI–stained nuclei (blue) in the ascending aorta from 20-week-old *Fbn1^C1041G/+^* and *Fbn1*^+/+^ mice (n=6). (**H**) Representative immunoblot analysis of Gfpt2, Uap1, O-GlcNac proteins in extracts from primary murine VSMCs transduced with *shFbn1* or *shScr (ShControl)* for 5 days (n=3). Actin was used as loading control. Data are mean±SEM. Statistical significance was assessed by Student *t* test. \**P*<0.05, \*\**P*<0.01 vs *Fbn1*^+/+^ mice.

Remarkably, in murine MFS-aortas, there was a significant upregulation of mRNA levels of key enzymes involved in the HBP, including the inducible rate-limiting enzyme Glutamine-fructose-6-phosphate transaminase 2 (*Gfpt2*), as well as downstream enzymes such as *Uap1* (UDP-N-Acetylglucosamine Pyrophosphorylase 1) and *Gnpat1* (glucosamine-phosphate N-acetyltransferase 1) (Fig. 1B). The enzymes O-GlcNAc transferase (*Ogt*) and O-GlcNAcase (*Oga*) (Fig. 1B), which mediate the dynamic cycling of O-GlcNac on a wide array of cytosolic, nuclear, and mitochondrial proteins were upregulated (Fig. 1A,B)^33^. Our analysis also revealed an increase in Golgi enzymes involved in extracellular protein N-glycosylation, including Alpha-1,3-Mannosyl-Glycoprotein 2-Beta-N-Acetylglucosaminyltransferase (Mgat) enzymes; *Mgat1, Mgat2, Mgat4a*, and *Mgat5b*, in MFS-aortas (Fig. 1B). Furthermore, the transcripts of genes such as *Gne* (Glucosamine UDP-N-Acetyl-2-Epimerase/N-Acetylmannosamine Kinase), *B3gnt3* (UDP-GlcNAc:BetaGal Beta-1,3-N-Acetylglucosaminyltransferase 3), and *Has1/Has3* (hyaluronan synthase 1/3), involved in the biosynthesis of N-acetylneuraminic acid, poly-N-acetyllactosamine chains, Sialyl LewisX (CD15s), and hyaluronic acid respectively; and *Cd44* (Hyaluronan receptor), were increased in MFS-aortas (Fig. 1B). These results were further corroborated by qPCR analysis of aortic extracts from MFS-mice (Fig. 1C). These findings suggest that the upregulation of HBP and downstream enzymes may contribute to elevated levels of O-GlcNAc, N-glycosylation and GAGs in the aortas of MFS mice.

Additionally, we conducted an analysis of *Gfpt2* mRNA expression in aortas at different ages. *Gfpt2* mRNA levels exhibit a significant increase at the very onset of aortopathy, as early as 4 weeks old, in *Fbn1^C1041G/+^* mice, even when the aortic phenotype is barely detectable^21^ (Fig. 1D). MFS-mice aortas show an increase of Gfpt2, Uap1 and O-GlcNAc levels by confocal immunostaining (Fig. 1E). MFS-mice aortas Immunoblot analysis confirmed elevated levels of O-GlcNAc and Gfpt2 levels in aortic samples from MFS-mice (Fig. 1F). To assess extracellular GAGs and glycoprotein levels, we used fluorescent Wheat Germ Agglutinin (WGA) that selectively binds to N-acetylglucosamine residues, predominantly present in the extracellular matrix, WGA staining is increased in MFS-mice^34^ (Fig. 1G).

In an effort to model MFS in vitro, we utilized lentiviral vectors to silence *Fbn1* expression in primary murine VSMCs^21^ (Suppl. Fig. 1A). Consistent with the transcriptomic data obtained from MFS mice, VSMCs with silenced Fbn1 (*shFbn1*) exhibited elevated protein levels of Gfpt2 and Uap1 (Fig. 1H) and *Gfpt2* mRNA levels (Suppl. Fig. 1A). Furthermore, *shFbn*1-VSMCs demonstrated increased levels of O-GlcNac and enhanced WGA staining, as determined by flow cytometry analysis (Fig. 1H; Suppl. Fig. 1B).

These findings suggest that *FBN1* deficiency induces abnormal HBP activity, leading to increased levels of O-GlcNac and WGA-staining in aortic tissue. These changes may contribute to the observed increase in GAGs and PGs observed in aortic medial degeneration in MFS aortas.

### Hexosamine Biosynthetic Pathway promotes aortic dilation and medial degeneration in wild type- and Marfan Syndrome-mice

Given that excessive GAGs-glycoprotein accumulation in aortic medial degeneration is a hallmark of TAAD^10,13^, we hypothesized that increased HBP activity might contribute to the PGs and GAGs accumulation, forming the ground-mucoid substance during TAAD development. To investigate this hypothesis, we treated *Fbn1^+/+^* and *Fbn1^C1041G/+^* mice with glucosamine (GlcN) to fuel HBP (Fig. 2A). Glucosamine is uptaken by the cells and undergoes transformation to glucosamine-6-phosphate though hexokinases and, ultimately converted to UDP-GlcNAc^23,35^ (Fig 1A).

**Figure 2:**
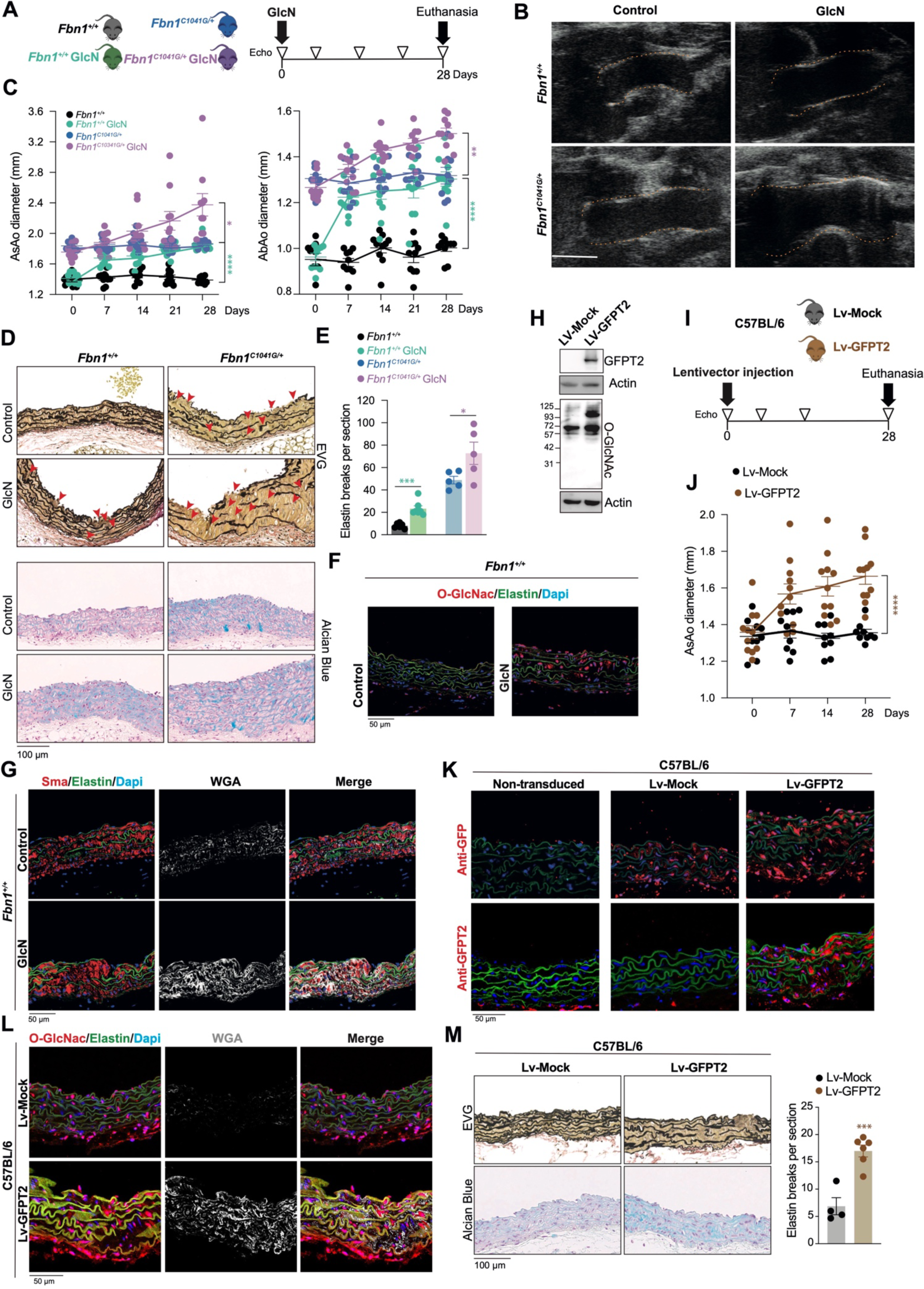
Boosting HBP induces aortic dilatation and medial degeneration. (**A**) Experimental design, panels A-E, 20-weeks old *Fbn1^C1041G/+^* and *Fbn1*^+/+^ mice were with or without glucosamine (GlcN in driking water for 28 days (**B**) Representative aortic ultrasound images after 28 days of control or GlcN treatment. Discontinuous red lines mark the lumen boundary, scale bar 1mm. (**C**) Evolution of maximal AsAo and AbAo diameter. (**D**) Representative histologic staining with EVG and Alcian blue in the AsAo and quantification of elastin breaks (**E**). **(F**) Representative confocal imaging of O-GlcNac proteins (red), elastin (green, autofluorescence), and DAPI–stained nuclei (blue) in the ascending aorta from *Fbn1*^+/+^ mice treated with or without GlcN for 28 days (n=6). **(G)** Representative confocal imaging of WGA (gray), Smooth muscle actin (Sma, red), elastin (green, autofluorescence), and DAPI–stained nuclei (blue) in the ascending aorta from *Fbn1*^+/+^ mice treated with/ without GlcN for 28 days (n=6). (**H**) Representative immunoblot analysis of Gfpt2 and O-GlcNac proteins in LV-Mock (control) or Lv-GFPT2 transduced primary murine VSMCs for 5 days (n=3). (**I**) Experimental design, panels J-M, 12-weeks old C57BL/6 wild-type mice were injected with LV-Mock or Lv-GFPT2 lentivectors for 28 days. (**J**) Evolution of maximal AsAo diameter after inoculation of LV-Mock (control) or Lv-GFPT2 lentiviral vectors. (**K**) Representative confocal imaging GFP or GFPT2 (red), elastin (green, autofluorescence), and DAPI-stained nuclei (blue) in ascending aortas from mice after 30 days of inoculation of LV-Mock (control) or Lv-GFPT2 lentivectors (n=6). (**L**) Representative confocal imaging of WGA (gray), O-GlcNac proteins (red), elastin (green, autofluorescence), and DAPI–stained nuclei (blue) in ascending aortas from mice after 30 days of inoculation of LV-Mock (control) or Lv-GFPT2 lentivectors (n=6). (**M**) Representative histologic staining with EVG and Alcian blue in the AsAo and quantification of elastin breaks in ascending aortas from mice after 30 days of inoculation of LV-Mock (control) or Lv-GFPT2 lentivectors (n=6). Data are mean±SEM. Statistical significance was assessed by 2-way repeated measurements ANOVA (**C,I**) or by 1-way ANOVA (**E**). \**P*<0.05, \*\**P*<0.01, \*\*\**P*<0.001, \*\*\*\**P*<0.0001 vs Control.

Treatment with glucosamine in drinking water for 28 days in wild-type mice (*Fbn1*^+/+^) led to a significant increase in aortic diameter (Fig. 2A-C). Notably, administration of glucosamine to *Fbn1^C1041G/+^* mice resulted in a further increase in aortic diameter compared to control *Fbn1^C1041G/+^* mice (Fig. 2A-C). The histological features of aortic medial degeneration were reproduced in *Fbn1*^+/+^ mice following glucosamine treatment and exacerbated aortic architectural medial abnormalities in *Fbn1^C1041G/+^*, such as elastin fiber fragmentation and glycan-matrix accumulation, as evidenced by Alcian Blue staining (Fig. 2D-E). Histological analysis also revealed that glucosamine increased HBP activity by O-GlcNac and WGA confocal stainings in aortic tissue from *Fbn1^+/+^* mice (Fig. 2F-G).

To elucidate the potential role of GFPT2 upregulation in the development of aortic aneurysm and aortic medial degeneration, we transduced primary murine VSMCs with GFPT2 encoding lentiviral vectors (LV-GFPT2). GFPT2 transduction resulted in an increase in O-GlcNac levels (Fig. 2H). LV-GFPT2 *In vivo* transduction in C57BL/6 wild-type mice (Fig. 2I), showed that increased GFPT2 expression led to an enlargement of aortic diameter (Fig. 2J). To further confirm lentiviral transduction and HBP activity, we performed confocal immunostaining of GFP, GFPT2 and O-GlcNac; WGA respectively (Fig 2K-L). GFPT2 transduction induce an increase in elastin breaks and Alcian Blue staining (Fig. 2M), indicating glycan-matrix accumulation.

These findings together suggest that excessive HBP activity resulting for GFPT2 upregulation contributes to the accumulation of mucoid substance in aortic medial degeneration during TAAD development and is detrimental to aortic homeostasis.

### Hexosamine Biosynthetic Pathway plays a critical role in the aortopathy of Marfan Syndrome

To investigate the therapeutic potential of HBP inhibitors in TAAD, we treated *Fbn1^C1041G/+^* - or *shFbn1*-VSMCs with DON (6-diazo-5-oxo-L-norleucine) for 48 hours. DON is an HBP inhibitor that blocks the enzymatic activity of GFPT. Treatment with DON efficiently reduced levels of O-GlcNac and WGA in *Fbn1^C1041G/+^* - or *shFbn1*-VSMCs, as evidenced by immunoblot and flow cytometry analysis (Fig. 3A; Suppl. Fig. 2A-B). Additionally, DON incubation resulted in decreased mRNA expression of classical MFS genes, such as *Tgb1*, *Fgf2*, Osteopontin (*Spp1*) and collagen1 (*Col1a1*) in *shFbn1*-VSMCs (Suppl. Fig. 2C).

**Figure 3:**
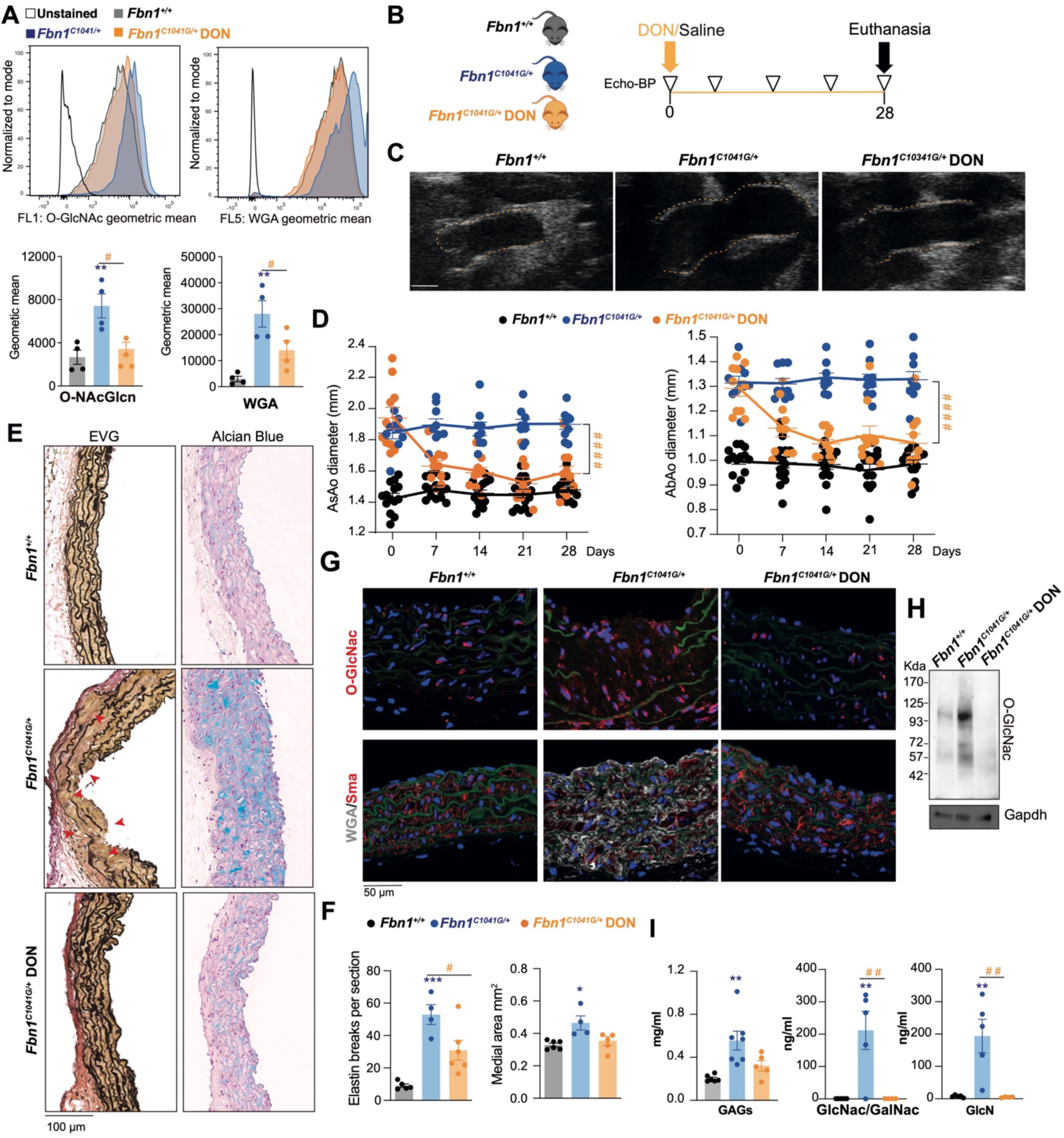
HBP inhibition by DON restores aortic dilatation and medial degeneration in MFS mice. (**A**) Representative flow cytometry histograms and statistical analysis of O-GlcNAc and WGA staining of primary VSMCs from *Fbn1^C1041G/+^* and *Fbn1*^+/+^ mice treated with or without DON for 24h. (**B**) Experimental design, panels B-J, 20/22-weeks old *Fbn1^C1041G/+^* and *Fbn1*^+/+^ mice were infused with minipumps with saline (control) or DON for 28 days. (**C**) Representative aortic ultrasound images after 28 days of control or DON treatment. Discontinuous red lines mark the lumen boundary, scale bar 1mm. (**D**) Evolution of maximal AsAo and AbAo diameter. (**E**) Representative histologic staining with EVG and Alcian blue in the AsAo and quantification of elastin breaks and aortic medial thickness (**F**) (n=6). **(G**) Representative confocal imaging of O-GlcNac proteins or SMA (red), WGA (gray), elastin (green, autofluorescence), DAPI–stained nuclei (blue) in the ascending aorta from *Fbn1*^+/+^ and *Fbn1^C1041G/+^* mice treated with/ without DON for 28 days (n=6). (**H**) Representative immunoblot analysis of O-GlcNac proteins in aortic extracts from *Fbn1^C1041G/+^* and *Fbn1*^+/+^ mice with saline or DON for 28 days (n=5). (**I**) GAGs serum levels, N-acetylGlucosamine / N-acetylGlucosamine and GlcN levels in sera from 20-week-old *Fbn1^C1041G/+^* and *Fbn1*^+/+^ mice with saline or DON for 28 days. Data are mean±SEM. Statistical significance was assessed by 1-way ANOVA (**A**, **F,J**) or 2-way repeated measurements ANOVA (**D**). \**P*<0.05, \*\**P*<0.01, \*\*\**P*<0.001, for *Fbn1^C1041G/+^* vs *Fbn1*^+/+^; #*P*<0.05, ##*P*<0.01, ####*P*<0.0001 for *Fbn1^C1041G/+^* DON vs *Fbn1^C1041G/+^*.

Next, we evaluated the therapeutic potential of DON in modulating HBP activity during the development of aneurysms in 20/22-weeks old MFS mice over a 28-day period was evaluated (Fig. 3B). Remarkably, after 28 days of treatment, DON normalized aortic dilation in *Fbn1^C1041G/+^* mice (Fig. 3C-D) and restored histological features of aortic degeneration in MFS, including medial thickening, elastic fiber fragmentation, and deposition of GAGs and PGs observed by Alcian blue staining (Fig. 3E). Furthermore, DON treatment for 28 days also resulted in reduced levels of O-GlcNac and WGA observed by immunoblot and confocal staining in aortic tissue (Fig. 3F-G). Notably, DON treatment in *Fbn1^C1041G/+^* mice, normalize GAGs and HBP-metabolites in serum, including N-AcetylGlucosamine/N-AcetylGalactosamine (GlcNac/ GalcNac), and glucosamine (GlcN) (Fig. 3H-I).

These data indicate that HBP inhibition decreases serum GAGs and HBP-metabolites, aortic glycosylation levels, thereby reducing aortic medial degeneration, improving aortic architecture, and restoring homeostasis in a MFS mouse model.

### Integrated Stress Response is activated by Hexosamine Biosynthetic Pathway in thoracic aortic aneurysm

The analysis of RNA-seq data from the aortas of 24-week-old MFS-mice revealed the we observed activation of Endoplasmic Reticulum Stress (p-value= 0.032, z-score=1.342). In the aortas of MFS-mice, the expression of genes associated with the Endoplasmic Reticulum-stress profile were increased, including *Atf4, Atf3, Atf6, Chop, Xbp1*, chaperones such as *Hspb1, Hspa5, Hspb7, Dnajc3*, and *prolyl 4-hydroxylase* (*P4hb*) (Fig. 4A). Endoplasmic Reticulum Stress occurs when unfolded or misfolded proteins accumulate in the Endoplasmic Reticulum, disrupting its normal function and leading to cellular perturbations^31^. To cope with Endoplasmic Reticulum-stress, cells activate the Integrated Stress Response (ISR), a complex signaling network. One critical component of the ISR is the phosphorylation of protein kinase PERK (PKR-like endoplasmic reticulum kinase), which phosphorylates the translation initiation factor eIF2α (eukaryotic initiation factor 2α). This phosphorylation event inhibits canonical protein synthesis while promoting the translation of specific mRNAs, including *ATF4* (Activating Transcription Factor 4). Through its downstream targets, ATF4 coordinates cellular adaptations aimed at restoring Endoplasmic Reticulum homeostasis and facilitating cell survival in the face of stress^31^. It has been described that the activation of the HBP through increased GFPT activity led to Endoplasmic Reticulum-stress, triggering the phosphorylation of both PERK and eIF2α, as well as downstream ATF4 translation, thereby identifying the HBP as a modulator of ISR^29^.

**Figure 4:**
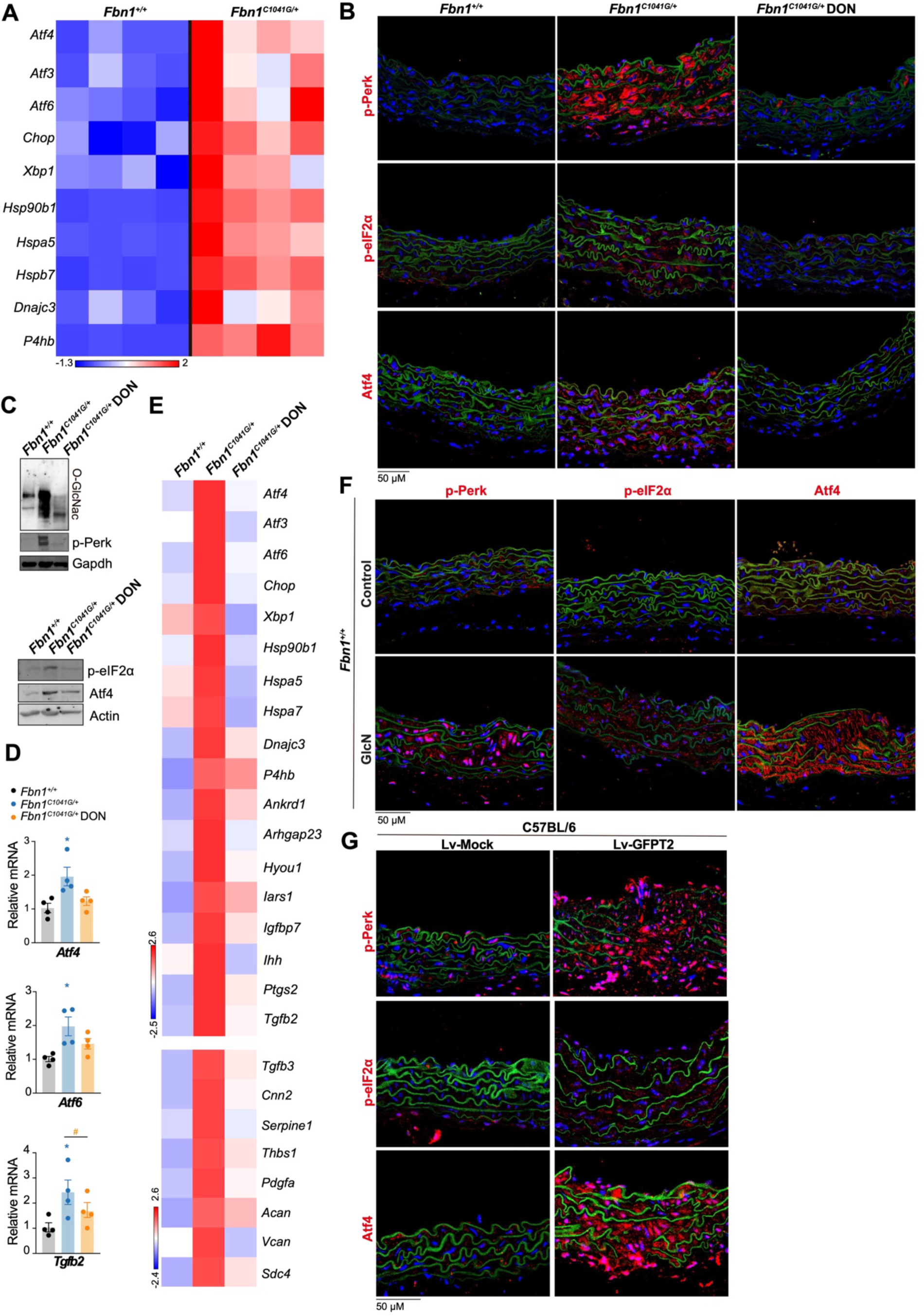
ISR is activated by HBP in aortic tissue. (**A**) Gene expression heatmap of ISR-related genes from RNA-sequencing analysis of aortic medial tissue from 24-weeks old *Fbn1*^+/+^ mice (n=4) and from *Fbn1^C1041G/+^* mice (n=4). (**B**) Representative confocal imaging p-Perk, p-eIF2*α* or atf4(red); elastin (green, autofluorescence), and DAPI–stained nuclei (blue) in ascending aortas from 20-week-old *Fbn1^C1041G/+^* and *Fbn1*^+/+^ mice infused with saline or DON for 28 days (n=5). (**C**) Representative immunoblot analysis of O-GlcNac proteins, p-Perk P-eIF2a or Atf4 in aortic extracts from 20-week-old *Fbn1^C1041G/+^* and *Fbn1*^+/+^ mice infused with saline or DON for 28 days (n=5). (**E**) Gene expression heatmap of ISR- and extracellular matrix-related genes from RNA-sequencing analysis of aortic medial tissue from 20 weeks old *Fbn1*^+/+^ mice (n=4) and from *Fbn1^C1041G/+^* mice infused with saline or DON for 28 days (n=4). (**F**) Representative confocal imaging p-Perk, p-eIF2*α* or Atf4(red); elastin (green, autofluorescence), and DAPI-stained nuclei (blue) in ascending aortas from 20/22 weeks-*Fbn1*^+/+^ mice treated with or without GlcN for 28 days (n=6). (**G**) Representative confocal imaging p-Perk, P-eIF2*α* or atf4(red); elastin (green, autofluorescence), and DAPI–stained nuclei (blue) in ascending aortas from 20-week-old *Fbn1*^+/+^ mice transduced with LV-Mock or LV-GPFT2 for 30 days (n=5). Data are mean±SEM. Statistical significance was assessed by 1-way ANOVA \**P*<0.05, for *Fbn1^C1041G/+^* vs *Fbn1*^+/+^; #*P*<0.05 for *Fbn1^C1041G/+^* DON vs *Fbn1^C1039G/+^*.

To determine the potential relationship between ISR and HBP in the vasculature, we treated *shFbn1*-VSMCs with DON for 48 hours. DON treatment led to a decrease in ISR signaling, evidenced by reduced phosphorylation of eIF2α and decreased protein and mRNA expression of *Atf4* and *Atf6* (Suppl. Fig. 3A-B). To assess whether the HBP activity induce ISR in MFS *in vivo*, we performed histological analysis of p-Perk, p-eIF2α and Atf4 staining in control and DON-treated for 28 days *Fbn1^C1041G/+^* mice. The results indicated that DON treatment reduces ISR in the aortas of MFS mice. (Fig. 4B). Moreover, we analyzed O-GlcNac, p-Perk, p-eIF2α and Atf4 protein levels by immunoblot (Fig. 4C) and mRNA levels of *Atf4* and *Atf6;* and also, Atf4-response gene *Tgfb2* (Fig. 4D). These data indicate that ISR is activated in MFS-mice and inhibited by DON treatment.

To characterize the molecular impact of DON treatment, we conducted RNA-sequencing analysis on aortas obtained from *Fbn1^+/+^* and *Fbn1^C1041G/+^* mice treated with or without DON for 28 days (n=4). Our results revealed that DON treatment led to decreased expression of genes associated with the ISR, including *Atf4* and Atf4-dependent genes such as *Chop, Xbp1, Hsp90b1*, *Ankrd3, Arhfap23, Hyou1, Iars1, Igfp7*, Cyclooxygenase-2 (*Ptgs2*), and *Tgfb2* (Fig. 4E). Remarkably, Ingenuity Pathway Analysis predicted a reduction in the TGFβ pathway activity in DON-treated mice, with a significant decrease in TGFβ upstream regulator activation Z-score (p-value= 2.57*10^−11^; activation Z-score= −9.913). Furthermore, DON treatment downregulated the expression of conventional MFS-affected genes, including transcripts involved in the TGFβ pathway and those encoding extracellular matrix-related proteins such as *Tgfb3, Cnn2, Serpine1, Thbs1, Pdgfa, Acan, Vcan,* and *Sdc4* (Fig. 4E).

To confirm that excessive HBP activity induces ISR in VSMCs, we analyzed eIF2α phosphorylation and Atf4 and Atf6 protein and mRNA gene expression in VSMCs transduced with a LV-GFPT2. This analysis revealed an activation of ISR (Suppl. Fig. 3C-D). Our *in vivo* data, revealed that augmenting HBP activity through glucosamine supplementation or LV-GFPT2 transduction in C57BL/6 wild-type mice for 28 days, leads to ISR signaling. This was evidenced by an increase in p-Perk, p-eIF2α, and Atf4 aortic immunostainings (Fig. 4F-G).

These findings indicate that excessive protein glycosylation induced by HBP activity triggers ISR signaling in aortas from wild type- and MFS-mice.

### Integrated Stress Response is involved in vascular smooth muscle cell dysfunction in Marfan syndrome

The ISR is a crucial cellular response pathway that regulates canonical protein translation rate and orchestrates a protective response, aiming to maintain cellular homeostasis and contribute to organismal fitness^31^. Initially, we examined Atf4 translation in MFS by transducing primary VSMCs from *Fbn1^C1041G/+^* and *Fbn1^+/+^* mice or *Fbn1*-silenced with a lentiviral vector carrying a reporter for Atf4 translation. This lentiviral vector contained a fluorescent protein, Scarlet, fused with the 5’ untranslated region of ATF4 (LV-ATF4Scarlet)^36^. Both *shFbn1*-VSMCs and VSMCs from *Fbn1^C1041G/+^* mice demonstrated an increase in Scarlet fluorescence compared to *shControl* or *Fbn1^+/+^* -VSMCs, indicating enhanced ATF4 translation ability in *Fbn1* deficient cells (Suppl. Fig. 4A; Fig. 5A). Furthermore, treatment of VSMCs-*Fbn1^C1041G/+^* with DON for 24 hours resulted in a decrease in Scarlet fluorescence compared to control VSMCs-*Fbn1^C1041G/+^*, pointing to the involvement of HBP in ATF4 translation (Fig. 5A).

**Figure 5:**
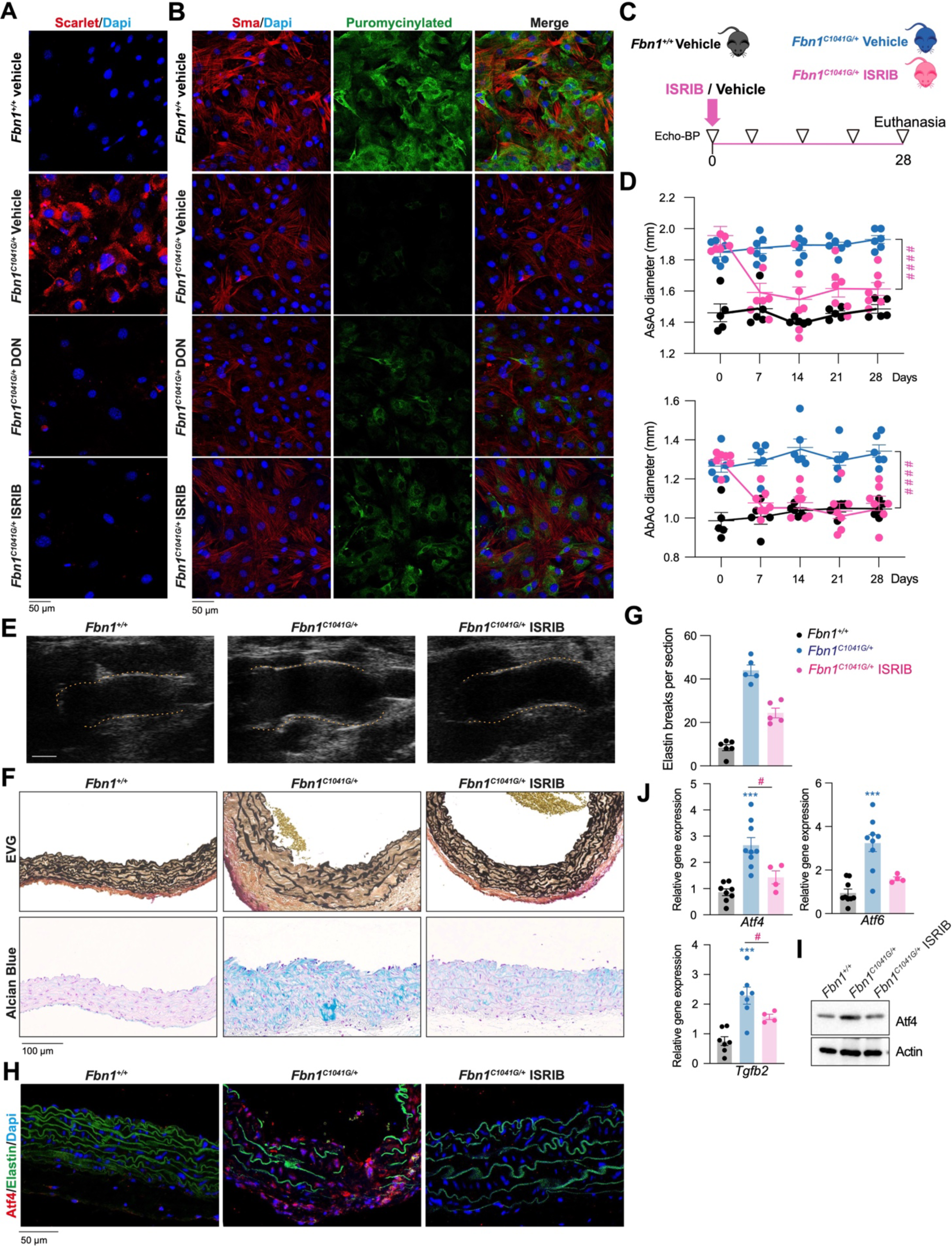
ISR contributes to VSMCs-dysfunction and aortic pathology in MFS-mice. (**A**) Representative confocal imaging of Scarlet fluorescence (red) in primary VSMCs (passage 2) from *Fbn1^C1041G/+^* and *Fbn1*^+/+^ mice transduced with LV-ATF4Scarlet after 3 days treated with Vehicle, DON or ISRIB for 48 hours. (**B**) Representative confocal imaging of puromycinylated proteins (green) and smooth muscle actin Sma, (Red) in primary VSMCs from *Fbn1^C1041G/+^* and *Fbn1*^+/+^ mice incubated with puromycin for 5 minutes and treated with vehicle, DON or ISRIB for 48 hours. (**C**) Experimental design, panels A-E, 20/22-weeks old *Fbn1^C1041G/+^* and *Fbn1*^+/+^ mice were infused with vehicle or ISRIB in minipumps for 28 days. (**D**) Evolution of maximal ascending (AsAo) and abdominal (AbAo) diameter. (**E**) Representative aortic ultrasound images after 28 days of control or ISRIB treatment. Discontinuous red lines mark the lumen boundary, scale bar 1mm. (**F**) Representative histologic staining with EVG and Alcian blue in the AsAo and quantification of elastin breaks after 28 days of vehicle or ISRIB infusion (**G**). (**H**) Representative confocal imaging of Atf4 (red), elastin (green, autofluorescence), and DAPI-stained nuclei (blue) in the ascending aorta from *Fbn1*^+/+^ and *Fbn1^C1039G/+^* mice treated with/ without ISRIB for 28 days (n=4). (**I**) Representative immunoblot analysis of Atf4 in aortic extracts from *Fbn1^C1041G/+^* and *Fbn1*^+/+^ mice with vehicle or ISRIB for 28 days (n=3). (**J**) Quantitative reverse transcription polymerase chain reaction analysis of *Atf4, Atf6 and Tgfb2* mRNA relative expression in aortic extracts from *Fbn1^C1041G/+^* and *Fbn1*^+/+^ mice infused with vehicle or ISRIB for 28 days. Data are mean±SEM Statistical significance was assessed by 2-way repeated measurements ANOVA (**B**) or 1-way ANOVA (**G,J**) or \*\**P*<0.01, \*\*\**P*<0.001, for *Fbn1^C1041G/+^* vehicle vs *Fbn1*^+/+^ vehicle; #*P*<0.05, ##*P*<0.01, ####*P*<0.0001 for *Fbn1^C1041G/+^* ISRIB vs *Fbn1^C1041G/+^* Vehicle.

Next, we explored the role of ISR in MFS. Initially, we assessed the efficacy of ISR inhibitor (ISRIB) *in vitro*. ISRIB is a small molecule known to reverse the effects of p-eIF2α and has been investigated in various mouse models, including Alzheimer’s disease, traumatic brain injury, and dementia^31,37^. Treatment with ISRIB for 24 hours in *shFbn1*-VSMCs resulted in reduced levels of *Atf4* and *Atf6* mRNA (Suppl. Fig. 4B). Additionally, ISRIB treatment for 24 hours in VSMCs-*Fbn1^C1041G/+^* transduced with LV-ATF4Scarlet led to decreased Scarlet fluorescence, indicating the involvement of ISR in ATF4 translation in *Fbn1^C1041G/+^* -VSMCs (Fig. 5A). The ISR decreases canonical translation rate through the phosphorylation of eIF2α. To further investigate the translation rate in MFS, we employed puromycin, an antibiotic that incorporates into nascent proteins, allowing visualization of actively translating ribosomes^38,39^. After treating *Fbn1^C1041G/+^* or *Fbn1^+/+^* VSMCs with puromycin for 5 minutes, we analyzed the puromycylated proteins by immunostaining. *Fbn1^C1041G/+^* -VSMCs exhibited a significant decrease in puromycylated proteins compared to *Fbn1^+/+^* cells, indicating a lower rate of protein translation in *Fbn1* mutant cells (Fig. 5B). Conversely, treatment with DON and ISRIB for 24 hours in *Fbn1^C1041G/+^* - VSMCs increased puromycylated protein staining, indicating that HBP induces a decrease of translation rate by the activation of ISR in MFS (Fig. 5B).

Subsequently, we evaluated the possible role of ISR in MFS aortic pathology *in vivo* by infusing 20/22-weeks old *Fbn1^C1041G/+^* mice with ISRIB for 28 days (Fig. 5C). Remarkably, ISRIB-treatment led to a decrease in aortic diameter in *Fbn1^C1041G/+^* mice, restoring them to normal levels (Fig. 5D-E). Moreover, ISRIB treatment notably improved aortic histology, as evidenced by reduced elastic fiber fragmentation and diminished Alcian blue staining in MFS-mice (Fig. 5F-G). Additionally, ISRIB treatment led to a reduction in Atf4 protein levels as observed through immunostaining and immunoblot analysis (Fig. 5H-I) along with decreased expression of IRS-induced genes such as *Atf4, Atf6* and *Tgfb2* (Fig. 5J). These data suggest that maladaptive ISR-response mediates aortic pathology in TAAD associated with MFS.

### Hexosamine Biosynthetic Pathway and Integrated Stress Response axis in aortas from patients with Marfan Syndrome

Next, we pursued to determine whether the involvement of HBP and ISR extends to patients with MFS. Histologic analysis of O-GlcNac and WGA staining confirmed a significant increase in HBP activity in human aortic MFS samples (Fig. 6A,B). Furthermore, in two distinct patient cohorts, MFS patients exhibited elevated serum and plasma levels of GAGs compared to controls (Fig. 6C). Furthermore, immunostaining of GFPT2 in the medial layer of aortic sections from MFS patients showed a significant increase compared to sections from organ transplant donors (Fig. 6D,E). Notably, immunostaining of p-PERK, p-EIF2α, and ATF4 indicated a clear activation of ISR in medial aortic samples from MFS patients (Fig. 6D,E). Taken together, these findings support the notion that the HBP-ISR axis may serve as important molecular markers and mediators of aortic pathology in human TAAD. Consequently, the evaluation of HBP-ISR inhibitors for the treatment of TAAD is warranted (Fig. F, graphical abstract).

**Figure 6:**
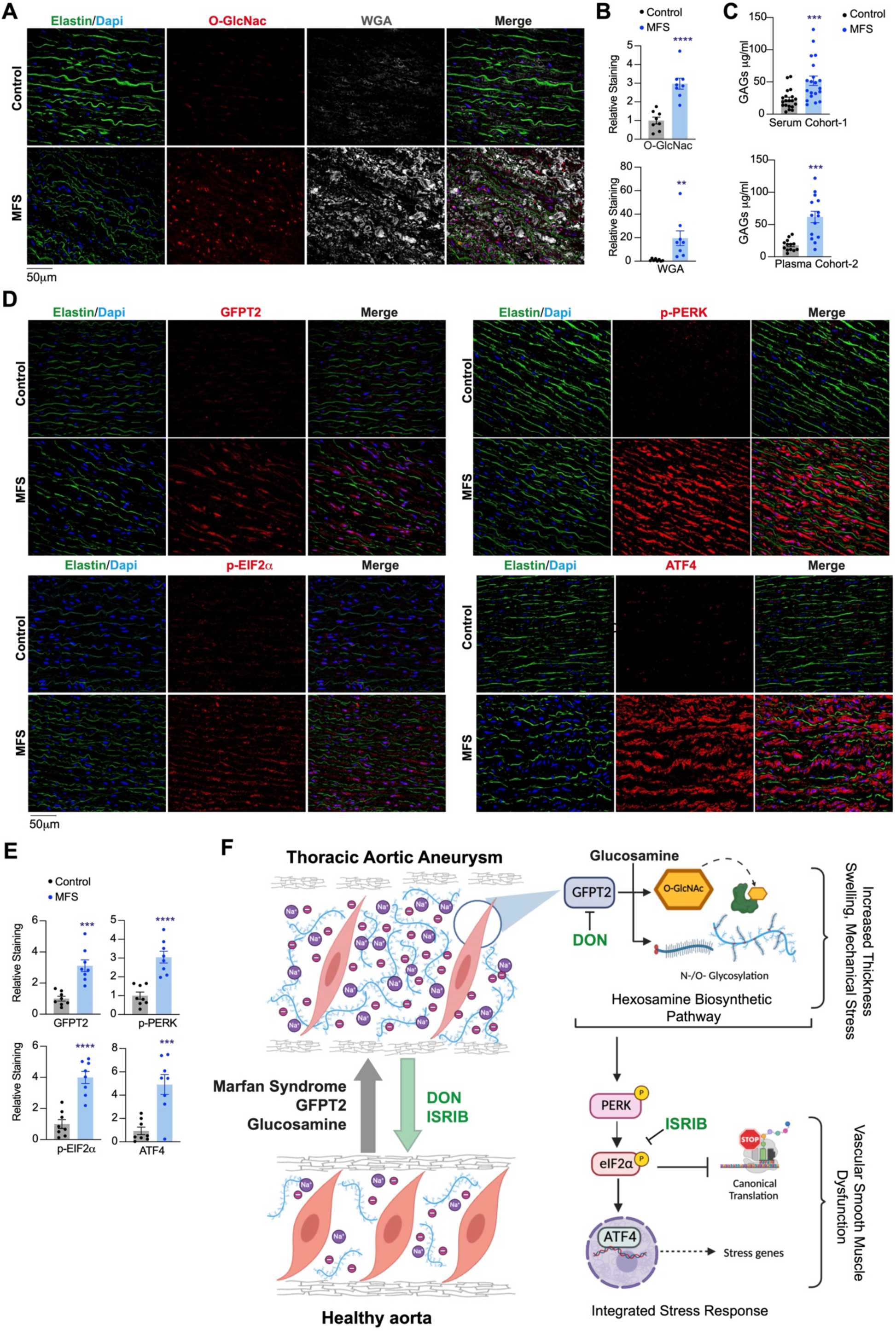
HBP-ISR axis is upregulated in MFS patients. (**A,D**) Representative confocal immunostainings and quantification (**B,E**) of O-GlcNac (A). (**C**) GAGs serum and plasmatic levels, in two different cohort of patiens. Representative confocal immunostainings and quantification; GFPT2, p-PERK, p-EIF2α, ATF4 (**E**) (red); WGA (A, gray), elastin (green, autofluorescence), and DAPI-stained nuclei (blue) in medial layer of aortic sections from control donors or patients with MFS (n=8; n=4 females, n=4 males; per group). Data are mean±SEM. (**G**) Proposed model depicting the differences between control and aortic pathology in MFS and subsequent cross-talk of the HBP-ISR axis. Statistical significance was assessed by Student *t* test. \**P*<0.05, \*\**P*<0.01, \*\*\**P*<0.01 \*\*\*\**P*<0.0001 vs Control. Falta la F que es el graphical abstract

## Discussion

Besides the high incidence and mortality risk associated with vascular complications, such as ruptured aneurysms or aortic dissection in patients with TAAD, therapeutic options to delay TAAD progression or the occurrence of dissection are limited, with none available to prevent it. Current pharmacological treatments for TAA primarily focus on blood pressure control using β-adrenergic blockers or the Ang II receptor-1 antagonist losartan, which can slow aortic dilation but do not prevent dissection^40,41^. Consequently, the management of aortic aneurysms often relies on surgical prophylactic repair, which carries significantly higher perioperative mortality and morbidity^42^. Thus, there is an urgent need to identify novel molecular mediators involved in heritable TAA pathophysiology in order to develop new pharmacological strategies^1,6^.

Our investigation revealed that the accumulation of glycoproteins, PGs and GAGs in TAAD is driven by excessive activity of HBP, a minor route in glucose metabolism responsible for protein glycosylation^23^. Although glucose flux through the HBP is minimal in homeostasis, it significantly increases in pathologies such as diabetes, cancer and heart hypertrophy^23^. The HBP acts as an integrator and sensor of key cellular metabolites, incorporating sugars (glucose), amino acids (glutamine), nucleotides (UTP), and fatty acids (Acetyl-CoA); hence, UDP-GlcNAc serves as a “metabolic sensor molecule”^23^ influencing various metabolic signaling pathways, including mTOR and AMPK, which contribute to pathophysiological alterations in aortic aneurysms^25,43,44^.

GFPT catalyzes the conversion of glutamine and fructose-6-phosphate into glucosamine-6-phosphate and glutamate. There are two mammalian paralogs of GFPT: GFPT1 and GFPT2, which share a high degree of sequence similarity, ranging from 75% to 80%. GFPT2 is expressed and induced in various cell types, including mouse embryonic stem cells, cardiomyocytes, macrophages, and neuronal cells. Notably, GFPT2 has been identified as a biomarker for poor prognosis malignancies such as breast, ovarian, colon, myosarcoma, and non-small cell lung cancer^23,24,45^. Unlike GFPT1, GFPT2 synthesizes glucosamine-6-phosphate more slowly but is not vulnerable to UDP-GlcNAc feedback inhibition. GFPT2 is inducible by NF-κB^46^ and TGFβ^47^ pathways, while GFPT1 is ubiquitously expressed^24^. Additionally, GFPT2 is involved in a TGFβ-pathway positive feedback loop by increasing AKT^35^ and TGFβ^48^ signaling pathways in cardiac fibrosis and hypertrophy. Importantly, we have demonstrated that both murine and human MFS show a marked increase in GFPT2 expression. Inhibition of HBP by DON leads to a reduction in the TGF-β-dependent genes within the aortas of MFS-mice, indicating the potential involvement in the positive feedback loop of the TGF-β pathway in MFS, thereby highlighting the potential of HBP as a therapeutic target (Fig. 6G). At present, specific targeted inhibitors for the GFPT1 or GFPT2 isoforms are unavailable. Considering the fundamental role of GFPT1-generated basal levels of HBP in cellular homeostasis and our observation of GFPT2 activity in mediating TAAD pathology, the development of novel therapeutics specifically targeting GFPT2 becomes crucial.

UDP-GlcNAc serves as a critical substrate for both O-GlcNAcylation and N-linked glycosylation processes. O-GlcNAcylation involves the addition of a single sugar, N-Acetylglucosamine, to serine or threonine residues on proteins, a process catalyzed by OGT^49^. This modification blocks further phosphorylation, hereafter, it is involved in many molecular signaling pathways. In contrast, O/N-linked glycosylation occurs predominantly in the Endoplasmic Reticulum and Golgi apparatus. Initially, O/N-linked glycan chains are attached to proteins in the Endoplasmic Reticulum, with subsequent branching and modification occurring in the Golgi, facilitated by enzymes such as MGATs. The resulting glycans are then secreted into the extracellular matrix. Moreover, UDP-GlcNAc serves as a donor substrate for the biosynthesis of GAGs such as hyaluronic acid, heparan keratan, and chondroitin sulfate, which are essential components of the extracellular matrix^50–52^. The dysregulation of HBP and the subsequent increase in enzymes such as HAS1/3, GNE, and PGs in MFS may contribute to the accumulation of ground substance in aortic medial degeneration. GAGs and PGs are highly negatively charged molecules that play important roles in maintaining the structural integrity of the extracellular matrix. Their negative charge allows them to maintain electroneutrality by attracting cations such as Na^+^, which in turn causes an influx of water molecules, leading to swelling of the aortic media. The accumulation of PGs and GAGs has a profoundly deleterious consequences in the aortic media, as it breaks the delicate micromechanical equilibrium that allows for proper function of the elastic elements and optimal mechanotransduction by the VSMCs^53–55^. The interstitial content of PGs/GAGs in aortic media is essential to maintain a specific level of wall hydration, enabling it to withstand cyclic compression while concurrently sustaining tension in the elastic elements and their connections with VSMCs^53^. This delicate mechanical equilibrium supports tensegrity and is pivotal for mechanotransduction in VSMCs. This elucidation emphasizes the importance of comprehending the structural alterations underlying aortic medial degeneration within the context of TAAD pathogenesis^56^. On the other hand, GAGs and PGs medial pooling disrupts the connections of VSMCs with their environment, favouring phenotypic switch and Anoïkis^57,58^. Inhibiting HBP by DON has been shown to impact the tumor microenvironment by reducing levels of hyaluronic acid and collagen, thereby facilitating anti-PD1 immunotherapy^59^. We observed reduced levels of GAGs and PGs by DON treatment This reduction in GAG and glycosylated protein accumulation has the potential to alleviate stiffness, swelling, elastin breaks and mechanical stress in the aorta, which may underlie the effects in attenuating TAAD development. Therefore, here, we have described for the first time how vascular metabolism is involved in the accumulation of GAGs and, consequently, in aortic medial degeneration. Further research is needed to fully elucidate the mechanisms by which GAGs and PGs contribute to aortic degeneration and how they interact with other components of the extracellular matrix to affect aortic biomechanics and integrity^20,56^.

Dynamic O-GlcNAc posttranscriptional modification of various cellular proteins plays a pivotal role in numerous biological processes, including cell proliferation, autophagy, apoptosis, stress response, epigenetics, metabolism, transcription, and protein trafficking^33^. Notably, important transcription factors implicated in TAAD pathogenesis, such as MYC, P53 and HIF1A^21^, are protected from degradation by O-GlcNac modification. Both types of glycosylation and enzymes involved are increased in mouse and human MFS-aortas. However, further investigation is warranted to elucidate the specific roles of O-GlcNAc and N-linked glycosylation in different aspects of TAAD development.

We have also observed that treatment with glucosamine increases glycosylation markers produced by the HBP in wild-type mice to levels similar to those occurring in MFS-mice aortas. Furthermore, glucosamine exacerbates aortic medial degeneration in MFS-mice. Glucosamine treatment has been shown to induce Endoplasmic Reticulum-stress and consequently activate the ISR in the context of diabetic atherogenesis^60^. Additionally, glucosamine is currently used as a dietary supplement to alleviate inflammation in osteoarthritis and joint pain in elderly patients, though its benefits remain unclear^61^. Given that many MFS patients suffer from joint pain, glucosamine supplements might be recommended. However, despite our experiments utilizing higher doses than those typically used in glucosamine supplements, we cannot disregard the potential harm of glucosamine based dietary supplements to patients at risk of vascular disorders. Further investigation to clarify this possibility is warranted.

Vascular aging is noticeable by distinct morphological changes, including arterial medial degeneration evidenced by heightened Alcian blue staining, medial calcification, and disarray in elastin fibers^62,63^. Indeed, it has been described that HBP and O-GlcNac are increased in aortas, heart, skeletal muscle and brain from aged rats^64^. Furthermore, observations in cardiac aging reveal an accumulation of GAGs due to HBP-increase^65^. Previous studies relate senescence in MFS-mice aortas^21,66^. Therefore, the augmentation of HBP throughout the aging process might significantly contribute to vascular senescence and associated pathological features in MFS.

Our findings suggest that N-acetylhexosamines and GAGs levels are elevated in the blood of both mice and patients with MFS, suggesting their potential as molecular markers for TAAD disease. These elevated levels imply dysregulation in glycosylation pathways, likely contributing to the pathogenesis of TAAD. Further investigation is essential for identifying glycosylated proteins in blood and aortic tissue, as this may elucidate the two branches of the glycomics landscape within TAAD and their correlation with the stage of aortic degeneration. Such exploration holds promise for uncovering novel molecular pathways, biomarkers, and potential therapeutic targets.

It is known that the ISR pathway activates as an adaptive response to various stress situations through four different kinases which converge into a common route. The stressors which these kinases recognize and act upon are heme-group deficiency, viral infection, amino acid deprivation and, most notably, Endoplasmic Reticulum stress^67^. HBP exhibits critical crosstalk with the Endoplasmic Reticulum-stress and therefore, ISR pathways, playing a pivotal role in maintaining proteostasis, particularly in aging cells^23,28,65,68^. Elevated HBP activity has been shown to promote ISR activation by increasing the phosphorylation of key mediators such as PERK and eIF2α, canonical translation inhibition and consequently leading to ATF4 translation^29^. In the context of TAAD, there may be a chronic maladaptive ISR response, which could contribute to VSMC dysfunction and apoptosis. The precise mechanisms by which augmented HBP activity affects Endoplasmic Reticulum-stress/ISR remain to be fully elucidated. However, it is conceivable that increased HBP activity might impact Endoplasmic Reticulum function by altering protein turnover, reducing protein synthesis, or increasing the formation of toxic protein-hyperglycosylated aggregates. Further research is warranted to explore whether targeting Endoplasmic Reticulum/ISR stress could be an effective therapeutic approach not only for TAAD but also for other vascular disorders such as cerebral or abdominal aortic aneurysms^69,70^. Such investigations could provide valuable insights into the broader implications of the HBP-ISR axis in vascular pathologies and inform the development of novel therapeutic strategies.

## Material and Methods

### Animal Procedures

Marfan mouse model harboring a *Fbn1^C1041G/+^* allele (JAX stock #012885). Wild-type littermates (*Fbn1^+/+^*) served as controls for the Marfan mice. Genotyping of the mice was conducted according to the protocols established by Jackson Laboratories. All mice used in the study were of the C57/BL6(CRL) background and male. For pharmacological interventions, DON (Sigma-Aldrich), dissolved in saline at a concentration of 5 ng* kg^−1^min^−1^, and ISRIB, dissolved in 20% DMSO-80% PEG400 (MedChemExpress), 1mg* kg^−1^min^−1^ were infused using subcutaneous osmotic minipumps (Model 2004, Alzet Corp). For control mice, the vehicle (Saline or DMSO/PEG400) was infused by subcutaneous osmotic minipumps. Glucosamine, 5% w/v (Merk), was given in the drinking water. To obtain mouse serum, blood was extracted after sacrifice by CO_2_ inhalation using the cardiac puncture method, collected and centrifuged for 15 min at 13,000 × g. Animal procedures and experiments complied with all relevant ethical regulations, were accredited by the CNIC and IIS-FJD Ethics Committee and the Madrid regional authorities (ref. PROEX 80/16, PROEX 74.7/23, PROEX 094.8/21), and conformed to EU Directive 2010/63/EU and Recommendation 2007/526/EC regarding the protection of animals used for experimental and other scientific purposes, enacted in Spanish law under Real Decreto 1201/2005. Mouse health was daily assessed for signs of discomfort; weight loss; or changes in behavior, mobility and feeding, or drinking habits. Mice were housed in an animal facility under a 12h light/dark cycle at constant temperature and humidity and had access to standard rodent chow and water *ad libitum*.

### Ultrasound imaging

*In vivo* ultrasound imaging was performed to monitor aortic diameter in isoflurane-anesthetized mice (2% isoflurane). High-frequency ultrasound was conducted using a VEVO 2100 echography device (VisualSonics, Toronto, Canada) with a resolution of 30 microns. Maximal internal aortic diameters were measured at diastole utilizing VEVO 2100 software, version 1.5.0.

### Cell procedures

Primary mouse VSMCs were isolated and cultured as previously described^71^. Briefly, aortic media tissue was digested with a solution of collagenase and elastase (Worthington Biochem) until a single-cell suspension was obtained. All experiments with primary VSMCs were conducted during passages 1–4. Treatments included DON (5nM), ISRIB (1μM) or Puromycin (1μg/mL for 5 minutes). Lentiviral transduction was performed in VSMCs over 5 hours at a multiplicity of infection = 3. The medium was then replaced with fresh DMEM supplemented with 10% FBS, and cells were cultured for three additional days. The HEK-293T (CRL-1573) and Jurkat (Clone E6-1, TIB-152) cell lines, required for high-titer lentivirus production and lentivirus titration, respectively, were purchased from ATCC. Experiments with these cell lines were conducted during passages 5–10. All cells were tested negative for Mycoplasma contamination.

### Lentivirus production and transduction

The lentiviral vectors coding sequences for human-GFPT2-GFP and Mock-GFP were obtained from Origene (RC200519L2); pLVX-ATF4 mScarlet NLS (Addgene plasmid # 115969). Lentivectors expressing shRNA targeting murine *Fbn1*, as well as control shRNA, were purchased from Sigma-Aldrich. Pseudotyped lentiviruses (VSV-G) were generated by transient calcium phosphate transfection of HEK-293T cells and concentrated from culture supernatant by ultracentrifugation (2 hours at 128,000xg; Ultraclear Tubes; SW28 rotor and Optima L-100 XP Ultracentrifuge; Beckman). The viruses were suspended in cold sterile PBS and titrated by transduction of Jurkat cells for 48 hours. Transduction efficiency (GFP-expressing cells and puromycin-resistant cells) and cell death (propidium iodide staining) were quantified by flow cytometry. For *in vivo* transduction experiments, animals were anesthetized with 2% isoflurane. A virus solution (100μl, 5*10^9^ viable particles/ml in saline) was inoculated directly into the retro-orbital sinus. Transduction efficiency was analyzed in aortic samples using immunofluorescence for GFP and GFPT2.

### Real-time and quantitative PCR

Aortas were extracted following perfusion with 5 ml saline solution, and the adventitia layer was discarded. Tissue frozen in liquid nitrogen was homogenized using a cold mortar and an automatic bead homogenizer (MagNA Lyser, Roche). Total RNA was isolated using Trizol (Life Technologies), with 1 μg of total RNA first digested with DNAse and then reverse-transcribed using the Maxima First Strand cDNA Synthesis Kit (ThermoFisher). Quantitative PCR (qPCR) reactions were performed in triplicate using SYBR master mix (Takara), following the manufacturer’s guidelines. Probe specificity was examined through post-amplification melting-curve analysis, ensuring that only one melting temperature (Tm) peak was produced for each reaction. Primer sequences as follows:

*Gfpt2* Fw GTATGATTGGCCGACCCTGG, Rv ATGCTAGCCGGAGAGCTGAA

*Uap1* Fw GCAGTTCTTCTTCTAGCTGGTG, Rv ATTCCATTGTTCTGCCGCTGGTC

*Gnpat1* Fw CAAATTGTGGCTACAGCAACT, Rv GACATTCAAGGGTGATCTTGTAAC

*Mgat2* Fw GCCATTACAGAGAGGCCAAG, AGAAACTCCGAATGGTGGTG

*Ogt* Fw GCCCTGGGTCGCTTGGAAGA, Rv TGCGACAGCTCTGTCAAAAA

*Oga* Fw CGGGAATTCCAGTGGCTTCG, Rv AAGCCGGGTGAACATCCCCA

*Atf4* Fw *A*AACCTCATGGGTTCTCCAG, Rv *GGC ATGGTTTCCAGGTCATC*

*Atf6* Fw AAT TCT CAG CTG ATG GCT GT, Rv TGG AGG ATC CTG GTG TCC AT

*Tgb2* CCACCTCCCCTCCGAAAA, Rv AGACATCAAAGCGGACGATTC

*Gapdh* Fw TGACGTGCCGCCTGGAGAAA, Rv AGTGTAGCCAAGATGCCCTTCA.

The amount of target mRNA in samples was estimated using the 2−CT relative quantification method, with Gapdh utilized for normalization. Fold ratios were calculated relative to mRNA expression levels from controls.

### Library preparation and Illumina sequencing

Aortas were extracted following perfusion with cold saline solution, and the adventitia layer was discarded. RNA from libraries was prepared according to the instructions of the “NEBNext Ultra Directional RNA Library Prep kit for Illumina” (New England Biolabs), following the “Poly(A) mRNA Magnetic Isolation Module” protocol. The input yield of total RNA to start the protocol was >300 ng, quantified using an Agilent 2100 Bioanalyzer with an RNA 6000 Nano LabChip kit. The obtained libraries were validated and quantified using an Agilent 2100 Bioanalyzer with a DNA7500 LabChip kit, and an equimolecular pool of libraries was titrated by quantitative PCR using the “Kapa-SYBR FAST qPCR kit for LightCycler480” (Kapa BioSystems) and a reference standard for quantification. Following processing on the Illumina HiSeq 2500 instrument, FastQ files were generated containing nucleotide data and quality scores for each position. RNA-seq reads were mapped to the *Mus musculus* reference genome, GRCm38.p6, using Hisat2 v2.1.0 software. Reads were then preprocessed with SAMtools v1.7 to transform SAM files into BAM files, which were subsequently sorted. These files served as input for the HTSeq v0.6.1 package, producing a file containing mapped reads per gene for each sample, as defined by the *Mus musculus*, GRCm38.96 version, gff file. After obtaining HTSeq read counts, the Bioconductor RNA-Seq workflow was followed to detect expression differences of genes using the DESeq2 statistical package. Ingenuity Pathway Analysis (IPA) was employed to identify cellular biological functions, gene clusters, and upstream regulators.

### Flow Cytometry and immunofluorescence

For flow cytometry analyses, primary VSMCs (passages between p1-p2) were trypsinized, fixed, permeabilized and sequentially incubated with fluorescein-RL2 antibody (Novus Biologicals), 647-WGA (Invitrogen). Samples were assessed with a FACSCanto II cell analyzer (Becton Dickinson) using DiVA acquisition software and FlowJo V10 data analysis software. For immunofluorescence, cells were fixed, permeabilized and incubated with and SMA-cy3 (Sigma) and anti-puromycin647 ThermoFisher antibodies.

### Metabolomics and GAGs determination

50 µl of serum samples (stored at −80 °C) and 360 µl of ice-cold methanol were vortexed for 2 min, then shaken in an Eppendorf shaker (Thermomixer R) at 800 rpm, 10 °C for 30 min and centrifuged at 10 °C for 10 min at ∼14,000 × g. Supernatants (320 µl) were transferred to a clean tube and evaporated to dryness under a stream of nitrogen gas at 40°C. Dried samples were resuspended in 70 μL of acetonitrile-water (80:20, v/v). The residue was reconstituted in 70 μL acetonitrile-water (80:20, v/v). LC-MS/MS analysis was performed using an integrated system composed of Agilent 1260 Infinity II and Ultivo 6465 Triple Quadrupole LC/MS system, which was equipped with an Agilent Jet Stream ESI source and controlled by MassHunter Workstation from Agilent Technologies (Santa Clara, CA, USA). Chromatography separation was performed on an ACQUITY UPLC BEH C18 column (1.7 µm, 100 mm × 2.1 mm i.d., Waters) and thermostated at 30 °C. The injection volume for each sample was 3 µL. The mobile phases were (A) water containing 0.1% formic acid (v/v) at pH 3 (adjusted with ammonium hydroxide) and (B) acetonitrile containing 0.01% formic acid (v/v). The analytes were eluted using the following program: 0-4 min, linear gradient 80-40% B; 4-5 min, linear gradient 40-20% B; 5-9 min, isocratic 20% B; 9-10 min, linear gradient 20-80% B; 10-16 min, isocratic 80% B to re-equilibrate the column. Quantitation by multiple reaction monitoring (MRM) analysis was performed in positive ion mode by monitoring the ion transitions m/z 222.1→204.0, 222.1→137.9 and 222.1→186.0 for GLcNAc, and m/z 180.1→162.0 and 180.1→72.1 for Glc. The source and gas parameters for the mass spectrometer were set as follows: ion spray voltage 4.0 kV, gas temperature: 325 °C, gas flow 10 L/min, nebulizer pressure 40 psi, sheath gas temperature: 350 °C, sheath gas flow: 12 L/min.

For GAGs determination, we utilized the Total Glycosaminoglycans Assay Kit (ab289842, abcam). We employed 10 μl of mice serum or 30 μl of human plasma/serum. Prior to GAGs-Probe incubation, we measured the absorbance of the samples to establish a baseline, which was then subtracted from the absorbance obtained after GAGs-Probe incubation.

### Immunoblot

For Western blot (WB) analysis, cells were lysed at 4 °C in RIPA buffer containing protease and phosphatase inhibitors cocktail (Promega/Sigma). Proteins were separated by SDS-PAGE and transferred onto 0.45 µm pore size Immobilon PVDF membrane (Millipore). PVDF membranes were blocked with TBS-T (50 mM Tris, 150 mM NaCl, and 0.1% Tween-20) containing 5% (w/v) milk. Membranes were incubated with primary antibodies diluted 1/500 to 1/1000 followed by TBS-T washes and incubation with HRP-conjugated secondary antibodies (GE healthcare). The signal was visualized by enhanced chemiluminescence with Luminata Forte Western HRP Substrate (Millipore) and the LightBright3 imaging system. The following antibodies were used: anti-O-GlcNac RL2 monoclonal (Abcam), Anti-GFPT2 (Abcam), anti-UAP1 (NovusBiologicals), anti-pPERK (ThermoFisher), and anti-pEIF2α and ATF4 (Cell signaling), anti Actin-HRP (Abcam) and anti-GAPDH (Abcam).

### Aortic histology

After euthanization by CO2 inhalation, mice were perfused with saline. Aortas were then isolated and fixed in 10% formalin overnight at 4 °C. Paraffin cross-sections (5μm) from fixed organs were stained with Alcian blue or Verhoeff elastic–van Gieson (EVG), or they were used for immunohistochemistry or immunofluorescence. Elastic fibers were stained with a modified Verhoeff Van Gieson elastin stain kit (Sigma-Aldrich). Elastic lamina breaks, defined as interruptions in the elastic fibers, were counted in the entire medial layer of three consecutive cross-sections per mouse. For immunostaining, the deparaffinized sections were rehydrated, boiled to retrieve antigens (10 mM citrate buffer, Triton-x 0,05%, pH 6) and blocked for 45 min with 10% goat normal serum, 5% horse serum, Triton-x 0,05% and 2% BSA in PBS. Samples were incubated with the following antibodies for immunohistochemistry or immunofluorescence: monoclonal anti-SMA-Cy3 (1/1.000, C6198, Sigma), WGA-647 (1/2.000 Invitrogen), polyclonal anti-GFPT2 (1/300, Abcam), Chicken anti-GFP (1/200, Abcam), monoclonal anti-O-GlcNac (1/200, Abcam), polyclonal anti-p-PERK (1/300 ThermoFisher) and anti-pEIF2, ATF4 (1/50 Cell Signaling). Specificity was determined by substituting primary antibody with unrelated IgG (Santa Cruz). For immunofluorescence, secondary antibodies were Alexa-Fluor-647-conjugated goat anti-rabbit, anti-mouse or anti-chicken (Invitrogen). Sections were mounted with DAPI in Prolong mounting medium (Invitrogen). Images were acquired at 1024 × 1024 pixels, using a Zeiss-LSM800 and Leica SP5 microscopes with 40× oil-immersion objectives.

### Human samples

The study complied with all relevant ethical regulations and was approved by the Clinical Research Ethics Committee of Cantabria (ref. 27/2013), the Ethics Committee of Instituto de Salud Carlos III (CEI PI91_2018-v2-Enmienda_2019 and CEI PI 65_2023) and Hospital Fundación Jiménez Díaz (ref. HBP2023) conformed to the principles set out in the WMA Declaration of Helsinki and the Department of Health and Human Services Belmont Report. Ascending aorta samples used as controls were obtained anonymously from multiorgan transplant donors. During preparation of the heart for transplantation, excess AsAo tissue was trimmed and harvested for the study. Aortic samples and data from MFS patients included in this study were obtained during elective or emergency surgery for aortic root replacement and provided by the Hospital Universitari Vall d’Hebron Biobank (National Registry of Biobanks B.000018, PT20/00107). 8 Control and 8 MFS samples were obtained from 4 male and 4 female individuals aged between 27 and 52 years old. Tissues were immediately fixed, kept at room temperature for 48 h, and embedded in paraffin. Informed consent was obtained from all human participants or their families. Patient clinical data were retrieved while maintaining anonymity. Plasma and serum samples from two cohorts of patients were collected: 20 controls and MFS patients with aortic aneurysms from Hospital Puerta del Hierro; and plasma samples from 15 controls and 14 MFS patients from Hospital Fundación Jiménez Díaz. These samples were obtained and processed in accordance with standard operating procedures.

### Statistical analysis

Normality of the data was assessed using the Shapiro-Wilk test, while the equality of variance assumption was evaluated using the F-test. Differences between two groups were analyzed using unpaired Student’s t-test, t-test with Welch’s correction for unequal variances, or Mann–Whitney U test where appropriate. Experiments with three or more groups were analyzed by one-way, two-way, repeated-measurements two-way analysis of variance (ANOVA), followed by Newman’s post hoc test as necessary. For statistical analysis of RNA-seq data, p-values were corrected using the false discovery rate (FDR) method (FDR <0.05). GraphPad Prism software 9 was used for all other analyses. Statistical significance was denoted as *P < 0.05, **P < 0.01, ***P < 0.001, and ****P < 0.0001. Sample sizes were determined empirically based on previous experiences in calculating experimental variability, without the use of statistical methods to predetermine sample size. Outliers were identified and excluded using the ROUT method (Q value = 5%) provided with GraphPad Prism 9 before data analysis. The number of animals used is described in the corresponding figure legends, with all experiments conducted with at least four biological replicates. Experimental groups were balanced in terms of animal age, sex, and weight. Animals were genotyped before experiments and housed together, receiving identical treatment. Sex and age are indicated in the figure legends. Appropriate tests were chosen based on data distribution, with comparable variance between groups in experiments described throughout the manuscript. Randomization was not employed for allocating animals to experimental groups, and investigators were not blinded to group allocation during experiments or outcome assessments.

## Acknowledgments

The authors wish to thank the plasma donors, and the Hospital Universitario Puerta de Hierro Majadahonda (HUPHM)/Instituto de Investigación Sanitaria Puerta de Hierro-Segovia de Arana (IDIPHISA) Biobank (Carlos III Health Institute Biobanks and Biomodels Platform) for the human specimens used in this study. We want to particularly acknowledge the patients and the Hospital Universitari Vall d’Hebron Biobank (PT20/00107) integrated in the Platform ISCIII Biobanks and Biomodels for their collaboration. Moreover, we want to acknowledge the patients from SIMA (Spanish Marfan Association) for their collaboration. pLVX-ATF4 mScarlet NLS was a gift from David Andrews (Addgene plasmid # 115969; http://n2t.net/addgene:115969; RRID: Addgene_115969). We thank the CNIC histology facility and animal imaging facility: A.V. Alonso and L. Flores for technical support. We thank to Citometry Unit from CBMSO. We thank the Conchita Rábago Foundation for supporting A.R-O. We thank the IIS-FJD animal facility and María del Mar González García-Parreño for technical support. We thank B. Ibañez for reagments, C. Ayuso for critical reading of manuscript and advice.

## Funding

A.R-O is supported by the Conchita-Rábago Foundation 2024 grant. J.O. is supported by a Ramón y Cajal contract (RYC2021-033343-I), and grant from Spanish Science Ministry (PID2022-1377300A-100); granted by Fundación MERCK-FEDER “Investigación clínica en enfermedades raras 2022”, Marfan Spanish association (SIMA, www.Marfan.es) and by “V-Ayudas Muévete por los que no pueden 2021”. Research in M.M.’s lab was supported by European Research Council (ERC-2021-CoG 101044248-Let T Be), by Comunidad de Madrid (Spain) (Y2020/BIO-6350 NutriSION-CM synergy) and by Spanish Ministerio de Ciencia e Innovación (PID2022-141169OB-I00) grants. JL.M-V is supported by grant from Spanish Science Ministry (PID2022-136979OB-I00). LM.B-C is supported by Fondo de Investigaciones Sanitarias, Instituto de Salud Carlos III (ISCIII/FEDER PI22/00233). G.T-T and A.G are supported by Spanish Society of Cardiology (SEC/FEC-INV-CLI 20/015). O.L is supported by frant from Spanish Health Ministry (PI20/00923 Instituto Carlos III). JF.N is supported by grant from Spanish Health Ministry (PI21/00084 FIS) and a grant from Health Research Institute Valdecilla (INNVAL21/24). N.M-V. is supported by Spanish Health Ministry (PI21/01126). MJ F-G is supported by Spanish Health Ministry contract (FIS22/00140). JM.R has received funding from “La Caixa” Banking Foundation (HR18-00068); Spanish Ministerio de Ciencia e Innovación grant PID2021-122388OB-100 funded by MCIN/AEI/10.13039/501100011033; and Instituto de Salud Carlos III (CIBER-CV CB16/11/00264); Fundació La Marató TV3 grant 202334-31, and Spanish Ministerio de Ciencia e Innovación contract FPI (BES-2016-077649) to MJ.R.-R. The CBMSO is supported by Consejo Superior de Investigaciones Científicas and Universidad Autónoma de Madrid. CBMSO is a Severo Ochoa Centers of Excellence (grant CEX2021-001154-S) funded by MICIN/AEI/10.13039/501100011033.

## Author Contributions

JO. and M.M. designed and conceived the study; A.R-O. and J.O. performed most of the experiments, with contributions from I.S-J, C.Z., MJ.F-G, MJ.R-R, A.L-Z and S.M-M. C.S. and V.G. perform metabolomics. S.M-A. extracted and processed blood plasma samples from patients. A.G, G.T-T and J.F.N. provided human tissue samples. TR.V. and A.F provided human serum samples. N.M-B, L.B-C., JL.M-V and JM.R. provided experimental support and ideas for the project; J.O. wrote the manuscript with contributions from N.M.B, A.R-O, E.G.R and M.M. All authors read and approved the manuscript.

## Notes

### Competing Interest Statement

The authors have declared no competing interest.

