## Supplemental material for "Integrated Stress Response Triggered by Excessive Glycosylation Drives Thoracic Aortic aneurysm"

### Supplemental Figures and Figure Legends

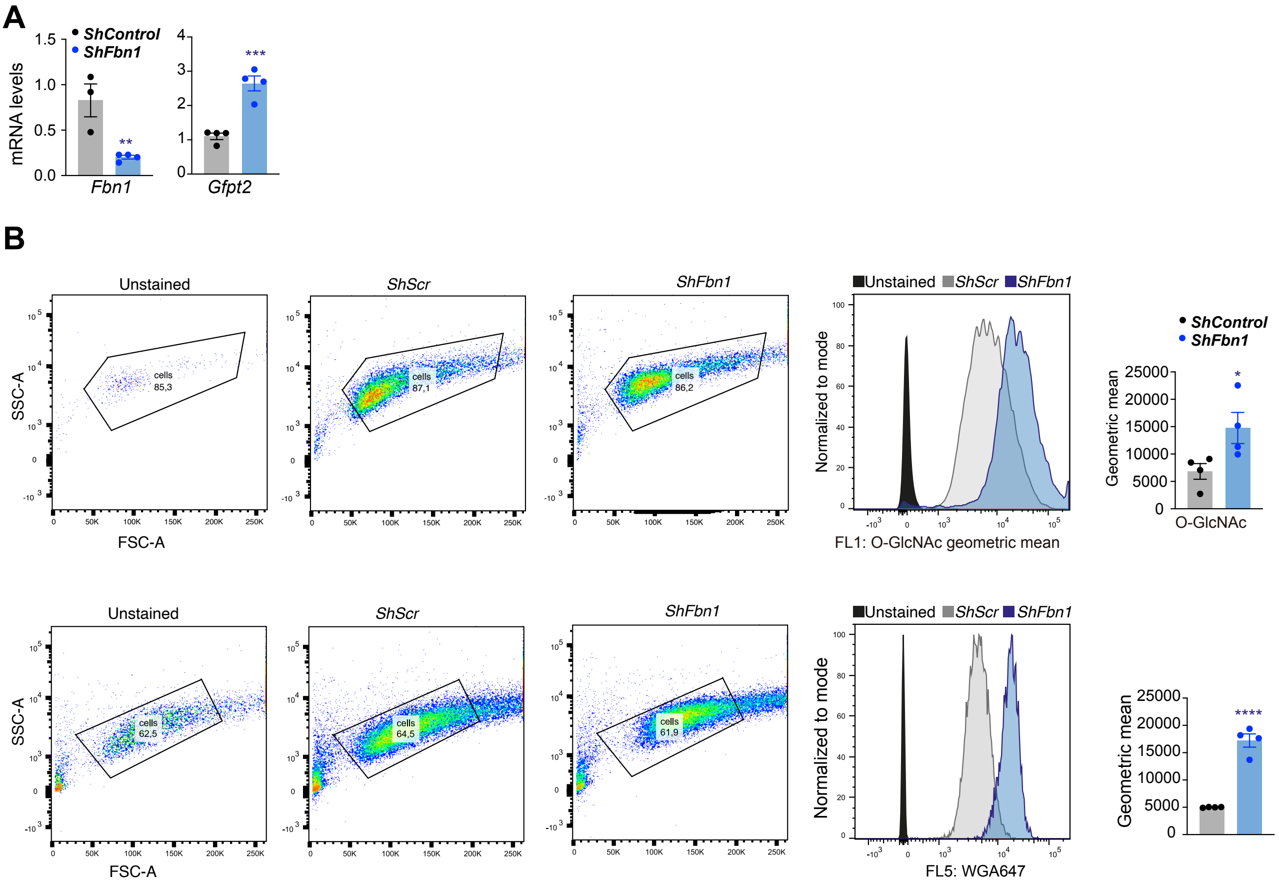


**Supplementary Figure 1: HBP increase in silenced Fbn1 VSMCs**

**(A)** Quantitative reverse transcription polymerase chain reaction analysis of *Fbn1* and *Gfpt2* in VSMCs after 5-days of shScr (shControl) or shFbn1 lentiviral vectors transduction. (**B**) Representative flow cytometry plots and quantification; VSMCs treated with or without DON for 24 hours, after 5-days of *shScr* (shControl) or *shFbn1* lentiviral vector transduction and were stained with O-GlcNac-Fluorescein/ WGA-647. Data are mean±SEM. Statistical significance was assessed by Student *t* test. **P*<0.05, ***P*<0.01, ****P*<0.001, *****P*<0.0001 vs *shScr*.

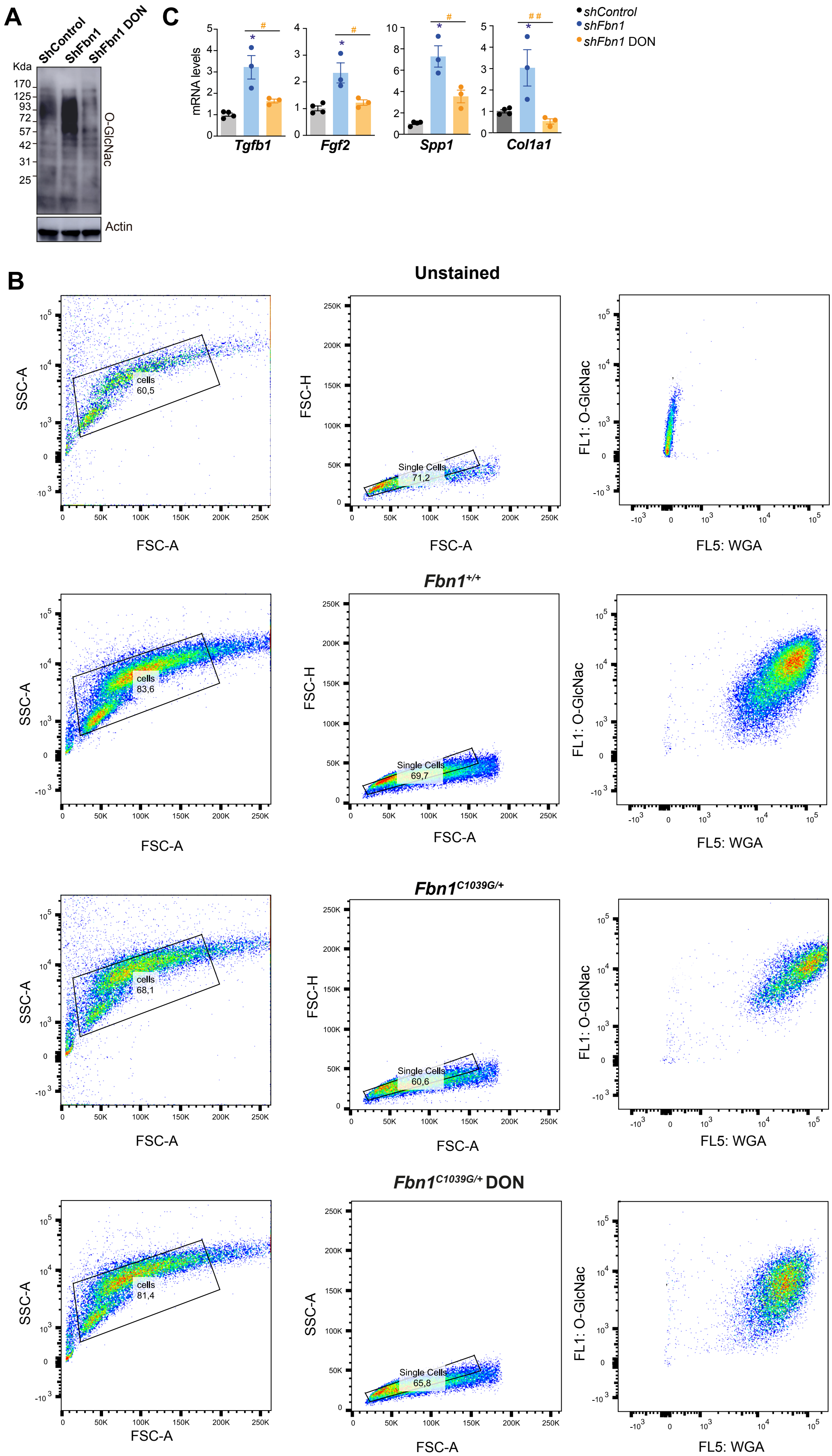
A
ShControl
ShFbn1
ShFbn1 DON
Kda
170
125
93
72
57
42
31
25
O-GlcNac
Actin
C
mRNA levels
Tgfb1
Fgf2
Spp1
Col1a1
shControl
shFbn1
shFbn1 DON
B
Unstained
Fbn1+/+
Fbn1C1039G/+
Fbn1C1039G/+ DON
SSC-A
FSC-A
FSC-H
cells
60,5
83,6
68,1
81,4
Single Cells
71,2
69,7
60,6
65,8
FL1: O-GlcNac
FL5: WGA

**Supplementary Figure 2: DON treatment in Fbn1-deficient VSMCs**

**(A)** Representative immunoblot analysis of O-GlcNac proteins in protein extracts from VSMCs treated with or without DON for 24 hours, after 5-days of *shScr* (shControl) or *shFbn1* lentiviral vector transduction, actin was used as loading control (n=3). (**B**) Representative flow cytometry plots shown in Fig. 4A; VSMCs from *Fbn1*$^{C1041G/+}$ and *Fbn1*$^{+/+}$ mice with or without DON for 24 hours, and were stained with O-GlcNac-Fluorescein/ WGA-647. (**C**) Quantitative reverse transcription polymerase chain reaction analysis of *Tgfb1, Fgf2, Spp1 and Col1a1* in VSMCs after 5-days of *shScr* (shControl) or *shFbn1* lentiviral vectors transduction and treated for 24h with or without DON. Data are mean±SEM. Statistical significance was assessed by 1-way ANOVA *$P$<0.05, **$P$<0.01, ***$P$<0.001, vs *shScr* ; #$P$<0.05, ##$P$<0.01, vs *shFbn1*.

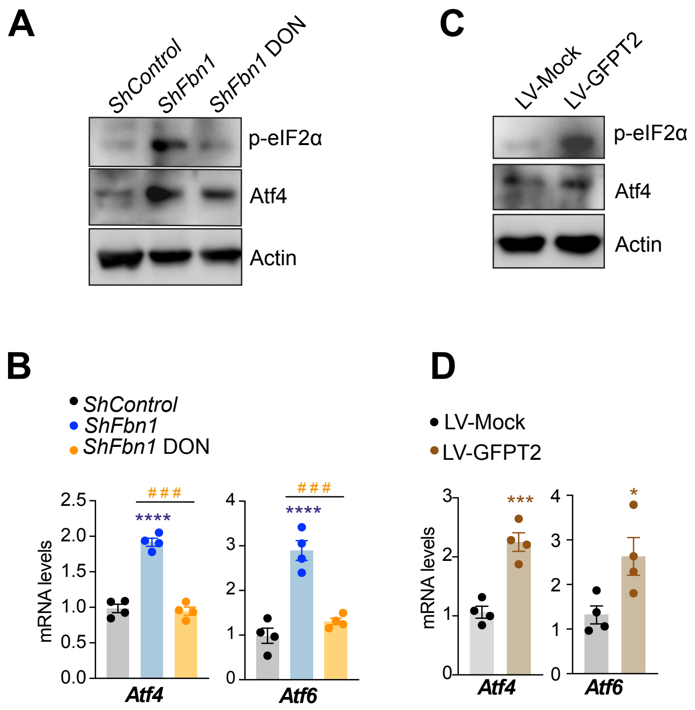


**Supplementary Figure 3: ISR is activated in Fbn1-deficient and LV-GFPT2 VSMCs**

**(A,C)** Representative immunoblot analysis of p-eIF2α and Atf4 proteins in protein extracts from VSMCs after 5-days of *shScr* (shControl) or *shFbn1* (**A**); and LV-Mock (Control) or LV-GFPT2 (**C**) lentiviral vector transduction , actin was used as loading control (n=3). **(B,D)** Quantitative reverse transcription polymerase chain reaction analysis of *Atf4 and Atf6* in VSMCs after 5-days of *shScr* (shControl) or *shFbn1* (**B**); and LV-Mock (Control) or LV-GFPT2 (**D**) lentiviral vectors transduction. Data are mean±SEM. Statistical significance was assessed by 1-way ANOVA (B) or student t-test (D) *$P$<0.05, ***$P$<0.001, vs *shScr* or LV-Mock; ##$P$<0.01, vs *shFbn1*.

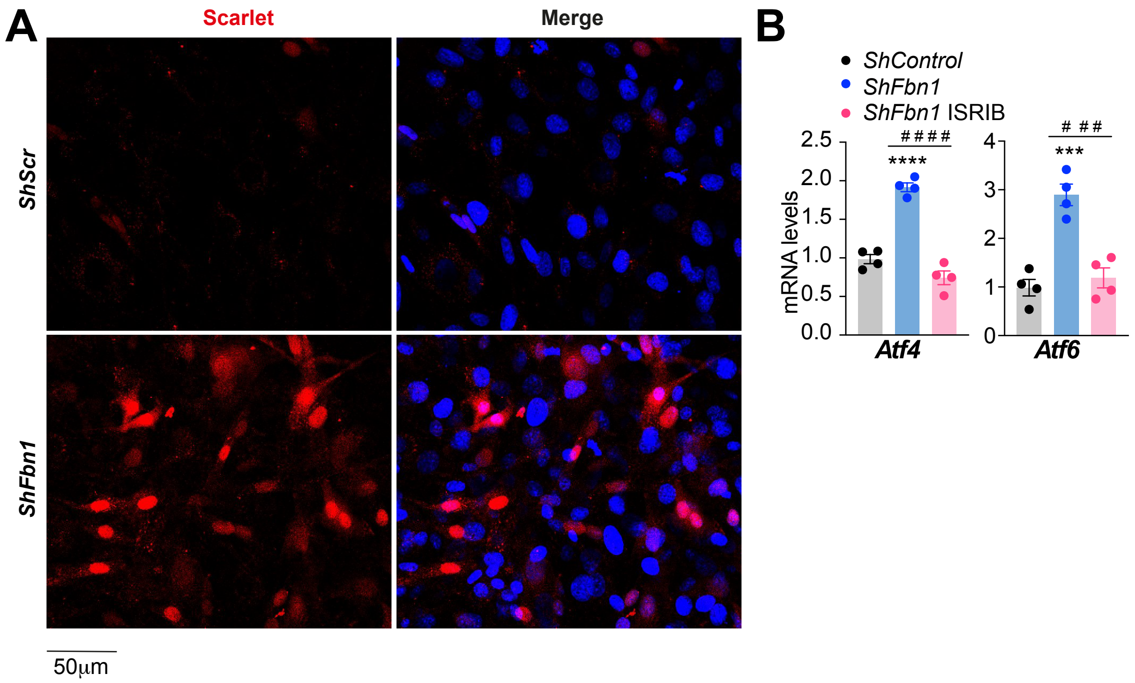


**Supplementary Figure 4: ISRIB treatment in Fbn1-deficient VSMCs**

**(A)** LV-ATF4Scarlet Lentiviral vector transduction control via scarlet (red) fluorescence in confocal imaging. (**B**) Quantitative reverse transcription polymerase chain reaction analysis of *Atf4 and Atf6* in VSMCs after 5-days of *shScr* (shControl) and *shFbn1* lentiviral vectors transduction after 5 days with or without ISRIB treatment for 24 hours. Statistical significance was assessed by 1-way ANOVA ****P*<0.001, *****P*<0.0001 vs *shScr* ; ###*P*<0.001, ####*P*<0.0001 vs *shFbn1*
